# Fractal Dimension and Lacunarity Measures of Glioma Subcomponents Provide a Quantitative Platform Discriminative of IDH Status: A Radiogenomics Approach in Gliomas

**DOI:** 10.1101/2023.12.28.573519

**Authors:** Neha Yadav, Ankit Mohanty, V Aswin, Navniet Mishrra, Vivek Tiwari

**Affiliations:** Indian Institute of Science Education and Research (IISER) Berhampur, Odisha, India

**Keywords:** Glioma, Fractal Dimension, Lacunarity, IDH, Machine Learning, Survival

## Abstract

**Background:** The presence of structural and geometric variations within gliomas, even among those with similar histologic grades, reflects the phenotypic heterogeneity unique to a genetic and epigenetic landscape. Whole glioma mass comprises of various subcomponents identified on MR imaging: enhancing, nonenhancing, necrosis, and edema fractions in varied fractions across patients. The geometry of whole tumor mass and the glioma subcomponents is highly irregular. Thereby, traditional Euclidean geometry is not suitable for quantifying the geometric dimensions. Here, we employ non-Euclidean geometric measurements: Fractal Dimension and lacunarity of the glioma subcomponents as a discriminator of IDH and MGMT status of gliomas.

**Methods:** Fractality and Lacunarity measurements were obtained using the tumor masks generated for enhancing, nonenhancing, and edema subcomponents from the preoperative T1, T1c, and T2-Flair MRI. Fractality and lacunarity measures of each subcomponent were evaluated between IDH mutant and wildtype gliomas. The fractality and lacunarity measures in IDH mutant and wildtype gliomas were further stratified for MGMT methylated and unmethylated gliomas. The fractality and lacunarities were trained and tested using supervised ML modeling as discriminators of IDH and MGMT status. Further, Cox Hazard estimations and the Kaplan-Meir investigations were performed to evaluate the impact of fractality and lacunarity measures of glioma subcomponents on the overall survival of the patients.

**Results:** IDH wildtype gliomas had ∼2-fold higher fractality for the enhancing subcomponent compared to IDH mutant enhancing subcomponent, while IDH mutant gliomas showed higher fractality for the nonenhancing subcomponent. Furthermore, the edema subcomponent did not differ for fractality or lacunarity measures between IDH mutant and wildtype gliomas. Fractal or lacunarity measures for either of the three subcomponents do not vary across MGMT methylated and unmethylated status with a given IDH mutant or wildtype gliomas. A combination of fractal measures of the enhancing and nonenhancing subcomponents together provided highly accurate and sensitive discrimination of IDH status using the supervised ML models. Moreover, fractality measure ≥ 0.69 for the enhancing subcomponent was associated with shortened patient survival: a fractal dimension value corresponding to that of IDH wild type gliomas. However, fractality and lacunarity estimates were not sensitive for discrimination of MGMT status.

**Conclusion:** Glioma structural heterogeneity measured as fractality and lacunarity using routine structural MRI measurements provide a noninvasive quantitative platform definitive of the molecular subtype of gliomas: IDH mutant *vs*. wildtype. Establishing fractality and/or lacunarity quantities as signatures of prognostic molecular events provides an avenue to bypass the need of biopsy/surgical interventions for decision-making, determining the molecular subtypes and overall clinical management of gliomas.

**Importance of the Study:** The non-Euclidean geometric measurements such as fractal dimension and lacunarity of enhancing, nonenhancing, and edema subcomponents are potentially unique quantitative metrics, discriminative of IDH status and patient survival. Fractality and Lacunarity estimates using the conventional structural MRI (T1w, T1C, T2, and T2F) provide an easy-to-use quantitative radiogenomics platform for improved clinical decisions, bypassing the need for immediate surgical interventions to ascertain prognostic molecular markers in gliomas, which is likely to improve overall clinical management and outcomes.

**Key Points:**

- Increased fractal dimensions of the enhancing subcomponents in IDH wildtype tumors, suggestive of highly irregular geometry, may potentially serve as a quantitative noninvasive determinant of IDH wildtype tumors.
- A combined fractal estimation of enhancing and nonenhancing subcomponents is the optimal and accurate discriminator of IDH mutant *vs*. wildtype.
- High fractal dimension of enhancing subcomponent and reduced fractality of nonenhancing subcomponent is predictive of shortened patient survival.

## Introduction

In 2016, WHO revised the glioma classification and introduced, for the first time, the molecular basis of glioma classification as IDH mutant and wildtype ^1^. Irrespective of the histologic grade, the gliomas harboring somatic mutations in the isocitrate dehydrogenase (IDH) gene confer better prognosis and improved survival compared to the IDH wildtype ^2,3^. In addition to the mutational events, a subset of gliomas harbors epigenetic methylation of proofreading enzyme O6-methylguanine-DNA methyltransferase (MGMT), while another subset has active MGMT^4^. Similar to the IDH mutants, gliomas with MGMT-methylation also have a good prognosis upon treatment with alkylating chemotherapy ^5^. Identification of mutational and epigenetic status requires tissue biopsy or surgical interventions. Given the complexity of surgery in the brain, clinical management of gliomas and other brain tumor patients is challenging ^6–9^. Given the high relevance of IDH and MGMT status in decision making for clinical management, it is imperative to have noninvasive quantitative clinically precise platforms definitive of IDH and MGMT status.

The structural and geometric heterogeneities observed across gliomas, even for similar histologic grades, are potentially a phenotypic manifestation of unique genetic and epigenetic events ^10^. Glioma mass is a heterogeneous bulk structure consisting of fractions of substructural components: enhancing (enhancing region of the tumor), nonenhancing, necrotic core, and edema (water deposition). Establishing the genome-phenome relationship by unraveling the geometry of enhancing, nonenhancing, necrosis, and edema subcomponents across gliomas harboring distinct molecular backgrounds; IDH mutant *vs.* IDH wildtype, MGMT methylated *vs.* MGMT unmethylated or a combination of IDH and MGMT events, is likely to establish a noninvasive quantitative method of subcomponent geometry definitive of glioma relevant molecular markers which may potentially help in improved clinical management and decision making.

Gliomas are complex and irregular structures that do not conform to traditional Euclidean geometry. Non-Euclidean geometric measurements, such as Fractal Dimension (FD or “Fractality”) and lacunarity, would better characterize the irregular and self-replicating patterns across gliomas of different molecular subtypes ^11–16^. FD serves as a metric for quantifying the complexity and irregularity of irregular shapes and structures. Gliomas possess inherent irregularity, rendering FD as an appropriate measure for assessing the extent of irregularity and its association with clinical outcome and molecular status ^17^. Another structural parameter, lacunarity, depicts the spatial distribution and arrangement of substructures inside the tumor mass. Lacunarity serves as a measure of the level of clustering or dispersion of substructural elements within a particular area of focus. Consequently, geometric uniqueness in the substructural phenotype of the glioma components may closely reflect the characteristics of the gliomas and associated genetic and epigenetic features.

Considering the structural heterogeneities even for the gliomas with the same histologic type and grades, here, we have performed a comprehensive investigation of FD and lacunarity in the glioma subcomponents using preoperative structural MRI for the glioma patients harboring IDH mutation and wildtype, and MGMT-methylated and unmethylated genetic and epigenetic events. In addition to the FD and lacunarity measures for the genetic subtypes, an ML-based radio-genomic platform discriminative of IDH and MGMT status was developed. Fractal measures of two subcomponents: enhancing and nonenhancing, together provides the most optimal feature discriminative of the IDH status.

## Methods

### Demographics

The study population included subjects with low grade glioma and glioblastoma. The imaging data utilized in the study were obtained from two repositories, namely TCGA-LGG (Low Grade Glioma) and TCGA-GBM (Glioblastoma Multiforme), provided by TCIA ^18^ (The Cancer Imaging Archive) of the National Cancer Institute. The TCIA is a publicly accessible database containing comprehensive radiological, genomic, and clinical data pertinent to various cancer types. The study considered only those subjects who had preoperative magnetic resonance imaging (MRI) scans consisting of T1-weighted, T2-weighted, T2w-fluid attenuated inversion recovery (T2-FLAIR), and contrast-enhanced T1-weighted (T1c) sequences together with manually verified tumor segmentation masks as well as age at diagnosis, gender, and IDH mutation status (biopsy confirmed).

### Tumor Segmentation

The tumor segmentation labels used in the study were generated using the GLISTR boost protocol ^19^. Consistent with the BraTS challenge, the generated labels were manually corrected, endorsed by an expert neuroradiologist, and made available for clinical analysis, model training, and evaluation ^20,21^. The segmentation labels later described the tumor subcomponents as enhancing, nonenhancing, and peritumoral edema.

### Fractal dimension and lacunarity calculation

An in-house pipeline in Python (v3.11) was established, implementing the 2D box counting function in the IMEA ^22^ to compute the FD in each axial slice for the whole tumor. The tumor region was extracted from each slice after binarization of the tumor mask and using contour detection and bounding rectangles as described in the 2D Fractal Dimension box-counting approach ^23^. After annotating the segmentations with the respective subcomponent labels, the box-counting method was applied to the defined bounding boxes for each tumor subcomponent to measure the FD across all the axial slices. The measures were computed, and the mean FD for each subcomponent was stored for further statistical investigations.

Lacunarity was estimated for each tumor subcomponent using the gliding box algorithm in combination with the binned probability distribution method ^24^. Within each box, the intensity distribution of pixel values was obtained as a histogram. To ensure the comparability between different box sizes, this histogram was then normalized using the effective breadth of the box. The calculation of lacunarity for each subcomponent was performed by dividing the sum of squares of pixel values, with each value weighted by its corresponding value in the probability density function. This division was done by the square of the probability density-weighted average of the pixels within the respective subcomponents. Subsequently, the mean lacunarity for each subcomponent was computed and recorded.

### Supervised Machine learning Modeling based on Fractality and Lacunarity Measures

To predict the tumor IDH status as IDH mutant and IDH wildtype, three supervised machine learning (ML) based algorithms: Support Vector Machine (SVM), Random Forest (RF), and k-Nearest Neighbors (KNN) were employed. The FD and lacunarity of the tumor subcomponents along with the IDH status were used to train the model wherein the FD and lacunarity estimates served as the dependent variable while the IDH status (mutant and wildtype) was used as the independent variable. The measures of fractality and lacunarity for the 142 subjects were divided into training and test data sets in an 80:20 ratio.

To ensure the reliability of the models, a 5-fold stratified cross-validation strategy was implemented. The data were partitioned into five folds such that each fold maintained a similar distribution of IDH mutant and wildtype subjects. The machine learning models were iterated five times such that four folds were used for training, and one fold was reserved as a test. A receiver operating characteristic (ROC) curve was obtained, and the Area Under Curve (AUC) was also calculated. A mean AUC ± standard deviation and confusion matrices were generated for all the 5-fold cross-validations. An average normalized confusion matrix was generated along with average prediction accuracies ± standard deviation to validate the overall performance of the models. A comprehensive examination was conducted, exploring all possible combinations of fractality and lacunarity of each tumor subcomponent to pinpoint the best FD or lacunarity combination of the subcomponents that served as the highly accurate and specific discriminator of the IDH status.

Similarly, ML procedures were extended to test the accuracy of discriminating MGMT status as methylated or unmethylated using either or both the FD and lacunarity estimates of each tumor subcomponent.

### Statistical analyses

All statistical analyses were performed using in-house scripts and available packages in Python (v3.11) and R (v4.1.2). The Mann-Whitney U test was used to evaluate the differences in the Fractal Dimension and lacunarity of tumor subcomponents across IDH mutant *vs.* wildtype. The cohort with confirmed IDH and MGMT status were further stratified in four groups based on the IDH and MGMT status; (A) IDH Mutant + MGMT methylated, (B) IDH Mutant + MGMT unmethylated, (C) IDH wildtype + MGMT methylated and (D) IDH wildtype + MGMT unmethylated. The fractal dimension and lacunarity across the four groups were compared using a non-parametric analysis of variance model (Kruskal-Wallis’ test). Adjusted p-values were used to estimate the differences between the fractal measures between the groups based on the molecular subtype combination. The threshold for statistical significance was set at p < 0.05 and was corrected for multiple comparisons using Bonferroni correction. Furthermore, the three supervised machine learning models; SVM, KNN, and RF, were implemented using the *scikit-learn* Python library ^25^. These models were trained using various combinations of FD and lacunarity of tumor subcomponents (enhancing, nonenhancing, and edema) to determine the accuracy and sensitivity of the models to discriminate IDH wildtype and IDH mutant status. Model performance was evaluated for each combination of tumor subcomponents using ROC-AUC, sensitivity, specificity, and normalized confusion matrices for accuracy. These matrices were used to evaluate the model’s precision for classifying subject’s IDH status. Similar statistical operations for ML models were conducted for MGMT status discrimination using FD and lacunarity measures.

To determine the optimal cutoff values for survival for Fractal Dimension and lacunarity within the three subcomponents, maximally selected rank statistics were applied ^26^ using the *maxstat* (v0.7.25) R package. The cutoff points were chosen such that it stratified for the groups having a size at least larger than 10% of the total cohort. Subsequently, the Kaplan-Meier survival analysis was performed in R (*survival* package ^27^) to estimate the overall survival between the two groups ; fractality or lacunarity cutoff values less than and more than the optimal cutoff for the respective subcomponents. The statistical significance of the difference in overall survival was estimated using the log-rank test.

Moreover, the univariate Cox Proportional Hazard (CPH) model was applied to explore the independent prognostic significance of the FD and lacunarity of each tumor subcomponent. Thereafter, the multivariate CPH model was established including all tumor subcomponents, for the Fractal Dimension and Lacunarity separately.

## Results

### Demographic and Clinical data

A total of 142 subjects (median age: 54 years) who cleared the inclusion criteria were investigated in the study. Based on the clinical features and demographic details; 62 subjects have low grade gliomas (Grade 2: 27; Grade 3: 35) and 97 subjects had glioblastoma (Table 1 and Supplementary Table S1). The mutation status of the IDH gene was biopsy confirmed for all the 142 subjects, while the methylation status of the MGMT gene was confirmed for 119 subjects. Overall survival (number of months and survival status) was available for the 141 subjects.

**Table 1.** Clinical Characteristics and Demographics.

| **Clinical Characteristics** | **Number (N)** |
| --- | --- |
| **Number of subjects** | 142 |
| **Age** |  |
| Median, years (range) | 54 (18 - 84) |
| **Gender** |  |
| Male (%) | 73 (51.4) |
| Female (%) | 69 (48.6) |
| **Glioma Grade** |  |
| Grade 2 | 26 |
| Grade 3 | 35 |
| Grade 4 | 81 |
| **Molecular Classification** |  |
| **IDH Status** |  |
| Wildtype | 87 |
| Mutant | 55 |
| **IDH & MGMT Status** |  |
| Mutant + Unmethylated | 6 |
| Mutant + Methylated | 49 |
| Wildtype + Unmethylated | 39 |
| Wildtype + Methylated | 25 |

### FD and Lacunarity across Tumor Subcomponents in IDH Mutant and Wildtype Gliomas

Fractal Dimension (FD) and lacunarity were determined for the glioma subcomponents using the subcomponent masks on MR images across IDH mutant and wildtype gliomas (Figure 1 A, B). The IDH wildtype gliomas presented with a significantly higher FD of the enhancing subcomponent compared to IDH mutant tumors (IDH Mutant: 0.53 ± 0.01, IDH wildtype: 1.28 ± 0.23; Difference = 0.75, 95% CI = 0.62 -0.92, p < 0.0001) (Figure 1C). Conversely, the IDH mutant gliomas had significantly higher FD for the nonenhancing subcomponent (IDH Mutant: 1.48 ± 0.18, IDH wildtype: 1.13 ± 0.27; Difference = 0.35, 95% CI = 0.27 - 0.46, p < 0.0001) (Figure 1D). However, edema subcomponent had no significant difference in FD between IDH mutant and wildtype gliomas (IDH Mutant: 1.93 ± 0.05, IDH wildtype: 1.92 ± 0.04; Difference = 0.01, 95% CI = 0.02 - 0.03, p = 0.54) (Figure 1E).

**Figure 1:**
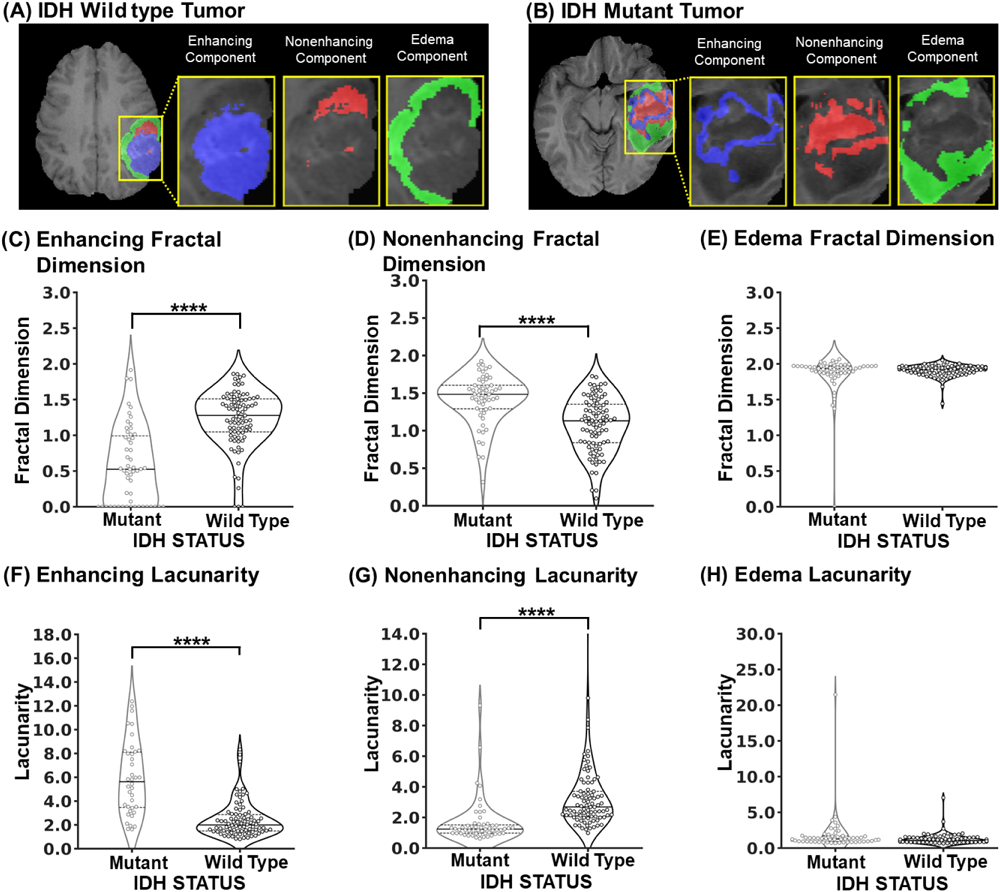
Glioma subcomponent mask and Fractality estimates between IDH mutant and wildtype gliomas. An axial slice of T1-weighted magnetic resonance image overlaid with the tumor mask of enhancing (blue), nonenhancing (red), and edema (green) subcomponents in (A) IDH wildtype and (B) IDH mutant glioma subjects. Violin plots depict the differences in the FD of (C) Enhancing, (D) Nonenhancing, and (E) Edema subcomponents between IDH mutant and IDH wildtype gliomas. (F-H) Differences in the lacunarity for Enhancing, Nonenhancing, and Edema subcomponents between IDH mutant and IDH wildtype tumors. Solid line shows the median and dotted line shows the interquartile range (IQR). The width of the violin plot shows the approximate frequency of data points in that region shows the kernel density depictive of probability of finding a data. * p<0.05, ** p<0.01, *** p<0.001, **** p< 0.0001, Mann-Whitney U test

The lacunarity of the enhancing component was significantly lower for the IDH wildtype gliomas compared to IDH mutant tumors (IDH Mutant: 5.63 ± 2.42, IDH wildtype: 2.00 ± 0.58; Difference = 3.6, 95% CI = 2.3 - 4.6, p < 0.0001) (Figure 1F) while the nonenhancing subcomponent presents with lower lacunarity (IDH Mutant: 1.24 ± 0.27, IDH wildtype: 2.68 ± 0.73; Difference = 1.44, 95% CI = 1.13-1.76, p < 0.0001) for the IDH mutant tumors compared to the IDH wildtype (Figure 1G). Similar to fractality measures, lacunarity across IDH mutant and wildtype was not significantly different (IDH Mutant: 1.27 ± 0.37, IDH wildtype: 1.07 ± 0.22; Difference = 0.19, 95% CI = 0.06 - 0.39, p = 0.05) (Figure 1H) between the two molecular subtypes of the tumors.

Furthermore, fractality comparisons across four groups based on MGMT status in IDH mutant and wildtype subjects (Table 1) did not exhibit significant differences between the fractal dimension and lacunarity for any of the tumor subcomponents (Figure 2). The, MGMT methylated gliomas with IDH mutation presented with lower fractal dimension for the enhancing subcomponent compared to the IDH wildtype MGMT methylated gliomas (IDH mutant + MGMT methylated median: 0.51 ± 0.50, IDH wildtype + MGMT methylated median: 1.21 ± 0.25, Difference: 0.70, 95% CI: 0.56 - 1.05, adjusted p < 0.001) (Figure 2A). For the fractal dimension of the nonenhancing subcomponent, MGMT methylated IDH mutant gliomas had increased fractal dimension compared to MGMT methylated IDH wildtype gliomas (IDH mutant + MGMT methylated median: 1.50 ± 0.15, IDH wildtype + MGMT methylated median: 1.14 ± 0.29, Difference: 0.36, 95% CI: 0.16 - 0.56, adjusted p < 0.001) (Figure 2C). For a given IDH status; fractal dimensions across any of the three subcomponents did not vary for methylated and unmethylated gliomas (Figure 2).

**Figure 2:**
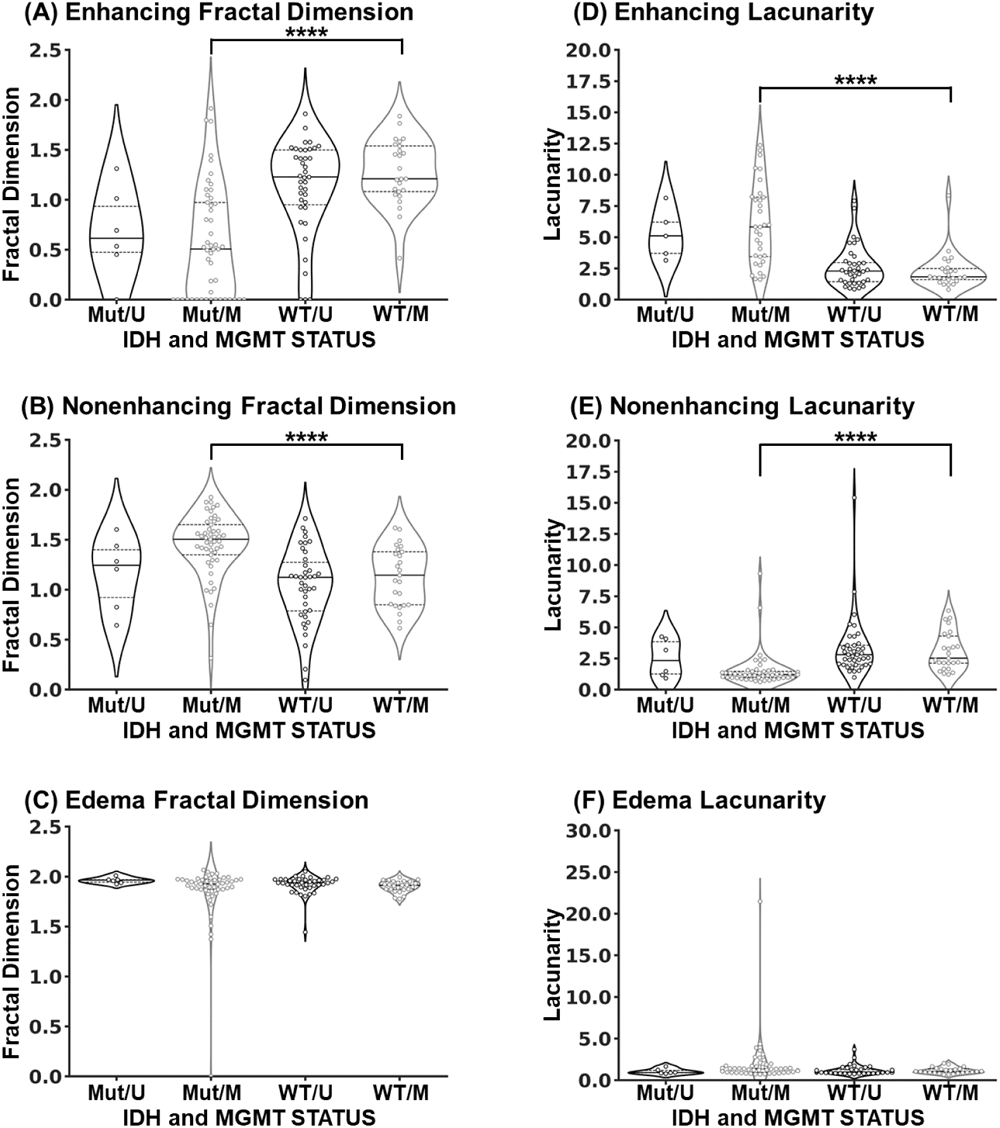
Fractality and Lacunarity of IDH mutant and wildtype gliomas stratified for MGMT status. Violin plots illustrate the differences in the (A) fractal dimension and (B) lacunarity of Enhancing; (C) fractal dimension and (D) lacunarity of Nonenhancing; and (E) fractal dimension and (F) lacunarity of Edema subcomponents for various possibilities of MGMT status in IDH mutant and wildtype gliomas. Solid line shows the median and dotted line shows the interquartile range (IQR). The width of the violin plot shows the approximate frequency of data points in that region shows the kernel density depictive of probability of finding a data. * p<0.05, ** p<0.01, *** p<0.001, **** p< 0.0001, Kruskal-Wallis test, post-hoc Dunn’s pairwise multiple comparison test. Mut, Mutant (IDH); M, Methylated (MGMT); WT, Wildtype (IDH); U, Unmethylated (MGMT)

The IDH mutant gliomas harboring MGMT methylation presented with higher lacunarity for the enhancing subcomponent (IDH mutant + MGMT methylated: 5.82 ± 2.38 vs IDH wildtype + MGMT methylated: 1.82 ± 0.46, Difference: 4.00, 95% CI: 2.26 - 5.32, adjusted p < 0.001) ( Figure 2B) and lower lacunarity for the nonenhancing subcomponent compared to IDH wildtype gliomas harboring MGMT methylated landscape (IDH mutant + MGMT methylated: 1.20 ± 0.24, IDH wildtype + MGMT methylated: 2.52 ± 0.90, Difference: 1.31, 95% CI: 0.90 – 2.20, adjusted p < 0.001) (Figure 2D). Noticeably, the glioma subsets stratified on basis of MGMT status across IDH mutant and wildtype did not exhibit significant difference of fractal dimension and lacunarity of the edema subcomponent (Figure 2E, F).

### Machine Learning based Classification and ROC analysis

The FD of each subcomponent in isolation, as well as in various combinations, was trained and tested for discriminative accuracy for IDH mutant and wildtype. The combination of the FD of enhancing and nonenhancing subcomponents provided the highest discriminatory accuracy across all three models (Figure 3C) with mean AUC values of 0.92 ± 0.08 using SVM (Accuracy = 0.86, SD = 0.08), 0.89 ± 0.08 using KNN (Accuracy = 0.87, SD = 0.08), and 0.88 ± 0.06 using RF (Accuracy = 0.83, SD = 0.08) (Figure 3B, C). The decision boundary of three supervised machine learning models depicts the discriminative accuracy as IDH mutant or wildtype, based on the distribution of FD (Figure 3A) and lacunarity (Supplementary Figure S1 A) of the enhancing and nonenhancing subcomponents.

**Figure 3:**
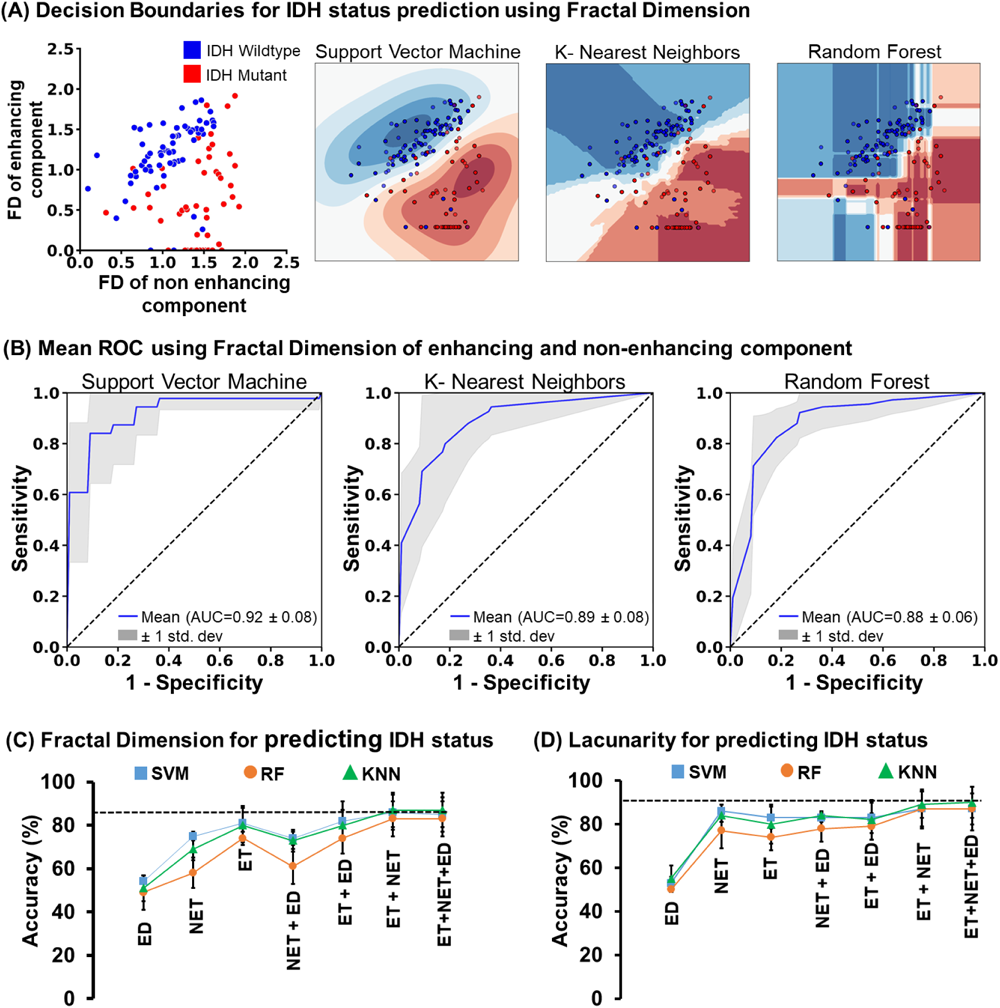
Fractality and Lacunarity based Machine learning models discriminative of IDH status. (A) Scatter plot depicting the distribution of IDH mutation status (Red - Mutant, Blue - Wildtype) using Fractal Dimension of enhancing and nonenhancing tumor subcomponents. Decision boundaries of three machine learning algorithms-SVM, RF, and KNN discriminating the IDH status. (B) The Receiver Operating Characteristic (ROC) curve of the three ML models: SVM, RF, and KNN, with mean AUC using the Fractal Dimension of enhancing and nonenhancing tumor subcomponents for distinguishing IDH molecular status. Accuracy of the three ML models: SVM (blue), RF (orange), and KNN (green) with various combinations of (C) Fractal Dimension and (D) Lacunarity of glioma subcomponents. ET, Enhancing tumor region; NET, Nonenhancing tumor region; ED, Edema tumor region

The FD of the enhancing subcomponent or nonenhancing subcomponent, individually, provides limited accuracy and sensitivity for IDH status discrimination (Figure 3C and Supplementary Figure S4 A). The FD of the edema subcomponent, individually or in combination with enhancing and nonenhancing subcomponents, did not have discriminative sensitivity (Figure 3C and Supplementary Figure S4).

Similarly, the lacunarity of each tumor subcomponent in all possible combinations was evaluated to discriminate the IDH status accurately. The combination of lacunarity of the enhancing and nonenhancing subcomponents provided high prediction accuracy of IDH status with mean AUC values of 0.89 ± 0.09 (Accuracy = 0.87, SD = 0.08), 0.90 ± 0.05 (Accuracy = 0.89, SD = 0.06), 0.92 ± 0.05 (Accuracy = 0.87, SD = 0.09) using SVM, KNN and RF models respectively (Figure 3D, Supplementary Figure S1.

The FD of the tumor subcomponents did not provide reliable accuracy for discrimination of MGMT status with the mean AUC of 0.70 ± 0.1 using SVM (Accuracy = 0.56, SD = 0.14), 0.69 ± 0.14 using KNN (Accuracy = 0.59, SD = 0.13) and 0.63 ± 0.06 using RF (Accuracy = 0.63, SD = 0.07) for the combination enhancing and nonenhancing tumor subcomponents (Supplementary Figure S2, S5 A). The lacunarity-based model for MGMT status also demonstrated lower discriminative sensitivity wherein a mean AUC of 0.73 ± 0.12 (Accuracy = 0.64, SD = 0.07), 0.71 ± 0.11 (Accuracy = 0.62, SD = 0.09), 0.70 ± 0.12 (Accuracy = 0.72, SD = 0.1) was obtained using the three models, respectively (Supplementary Figure S3, S5 B).

### Patient Survival and Hazard Analysis Using Cox Proportional Hazard (CPH) Model

A log rank statistics based estimation of a threshold of FD and lacunarity revealed a cutoff value of fractality and lacunarity for each tumor subcomponent such that it stratified the glioma subjects into two groups: below the cutoff and above the cutoff. The cutoff value of 0.69 for the fractality of the enhancing component, 1.2 for the nonenhancing component, and 1.8 for the edema subcomponent were used to evaluate the association between Fractality and overall survival (Figure 4). Fractal dimension more than the cutoff value of 0.69 for the enhancing subcomponent showed significantly shortened (*p < 0.0001 by the log rank test)* overall survival (OS = 17.1 months) compared to the subjects with FD < 0.69 (OS = 44.0 months) (Figure 4D). For the subjects with FD > 0.69 in the enhancing subcomponent, the hazard ratio for death was 3.9 (95% confidence interval, 1.9 - 8.2) (Table 2A).

**Figure 4:**
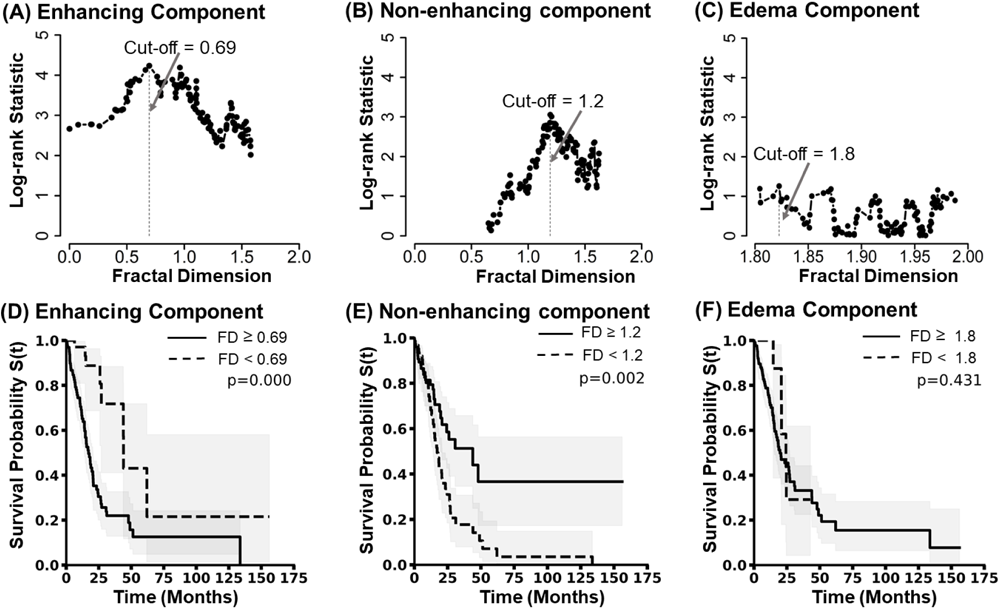
Fractality and Lacunarity based determination of Overall Patient Survival. Optimal cutoff values for the FD of (A) enhancing, (B) nonenhancing, and (C) edema subcomponents were determined using MaxStat and log-rank tests. Kaplan-Meier survival curves between the two groups formed using the optimal cutoff values of The Fractal Dimension of (D) enhancing, (E) nonenhancing, and (F) edema tumor subcomponents. The group with an FD cutoff greater than and equal to the cutoff is represented by the solid line and the group with an FD less than the cutoff value is represented by the dashed line.

**Table 2A.**
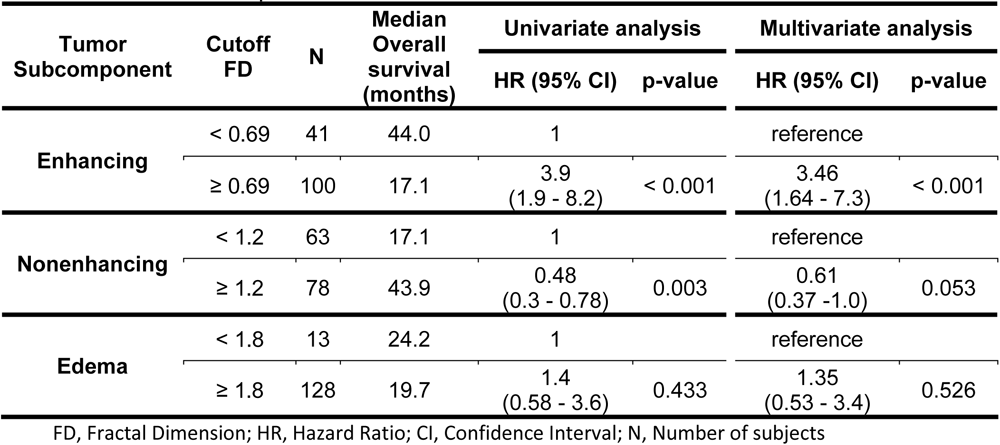
Univariate and multivariate Cox proportional hazard ratio for Fractal Dimension of tumor subcomponents.

| Tumor Subcomponent | Cutoff FD | N | Median Overall survival (months) | Univariate analysis |  | Multivariate analysis |  |
| --- | --- | --- | --- | --- | --- | --- | --- |
|  |  |  |  | HR (95% CI) | p-value | HR (95% CI) | p-value |
| Enhancing | < 0.69 | 41 | 44.0 | 1 |  | reference |  |
|  | ≥ 0.69 | 100 | 17.1 | 3.9<br>(1.9 - 8.2) | < 0.001 | 3.46<br>(1.64 - 7.3) | < 0.001 |
| Nonenhancing | < 1.2 | 63 | 17.1 | 1 |  | reference |  |
|  | ≥ 1.2 | 78 | 43.9 | 0.48<br>(0.3 - 0.78) | 0.003 | 0.61<br>(0.37 - 1.0) | 0.053 |
| Edema | < 1.8 | 13 | 24.2 | 1 |  | reference |  |
|  | ≥ 1.8 | 128 | 19.7 | 1.4<br>(0.58 - 3.6) | 0.433 | 1.35<br>(0.53 - 3.4) | 0.526 |

FD, Fractal Dimension; HR, Hazard Ratio; CI, Confidence Interval; N, Number of subjects

Kaplan–Meier estimates of overall survival in the two subgroups for the nonenhancing subcomponent (based on FD cutoff value = 1.2) were significantly different (p=0.002 by the log-rank test), wherein a FD value < 1.2 is associated with shortened survival (OS = 17.1 months) (Figure 4E). The hazard ratio for death was 0.48 (95 percent confidence interval, 0.30 - 0.78) for the subjects with a fractal dimension ≥1.2, corresponding to a 52% decrease in the risk of death (Table 2A). The interaction between the fractality of the edema subcomponent and overall survival (Figure 4F) was not statistically significant, according to the Cox proportional hazards model (p = 0.43) (Table 2A).

The log rank statistics revealed lacunarity cutoff values of 3.52 for the enhancing, 1.48 for nonenhancing, and 0.97 for edema subcomponents, stratifying the group in two groups (Supplementary Figure S7 A, B, and C). The cutoff value of lacunarity for each subcomponent, when subjected to Kaplan-Meier analysis for estimating the impact on overall survival, revealed that a lacunarity less than 3.52 in the enhancing subcomponent showed significantly shorter overall survival (OS = 15.3 months, p < 0.001 by the log rank test) compared to subjects with lacunarity > 3.52 (OS = 43.9 months) (Supplementary Figure S7 D). For the nonenhancing component, lacunarity greater than the cutoff of 1.48 was strongly linked to shorter overall survival (OS = 15.3 months, p < 0.001) (Supplementary Figure S7 E) with a significantly higher hazard ratio of mortality (HR = 4.1, 95% CI = 1.9 - 8.6) (Table 2B).

**Table 2B.** Univariate and multivariate Cox proportional hazard ratio for Lacunarity of tumor subcomponents.

| Tumor Subcomponent | Cutoff Lacunarity | N | Median Overall survival (months) | Univariate analysis |  | Multivariate analysis |  |
| --- | --- | --- | --- | --- | --- | --- | --- |
|  |  |  |  | HR (95% CI) | p-value | HR (95% CI) | p-value |
| Enhancing | < 3.52 | 81 | 15.3 | 1 |  | 1 |  |
|  | ≥ 3.52 | 39 | 43.9 | 0.3<br>(0.16 - 0.57) | < 0.001 | 0.47<br>(0.23 - 0.96) | 0.039 |
| Nonenhancing | < 1.48 | 33 | 62.0 | 1 |  | 1 |  |
|  | ≥ 1.48 | 87 | 15.3 | 4.1<br>(1.9 - 8.6) | < 0.001 | 2.54<br>(1.08 - 5.97) | 0.032 |
| Edema | < 0.97 | 48 | 15.0 | 1 |  | 1 |  |
|  | ≥ 0.97 | 72 | 24.5 | 0.52<br>(0.32 - 0.84) | 0.007 | 0.53<br>(0.32 - 0.86) | 0.01 |

HR, Hazard Ratio; CI, Confidence Interval; N, Number of subjects

Interestingly, from the Kaplan-Meier estimates, we observed that the group with lower lacunarity of edema component than the cutoff (0.97) had shorter survival than the group with higher lacunarity (*p = 0.006 by log rank test*) (Supplementary Figure S7 F). The latter group demonstrated a lower hazard ratio for mortality (0.52, 95% CI - 0.32 - 0.84) (Table 2B).

## Discussion

Even the gliomas that have similar histologic grade display diverse overall tumor shape and geometry. The structural and geometric heterogeneity of tumors may primarily arise from the highly irregular geometry of the tumor subcomponents, such as the enhancing, nonenhancing, necrotic, and edema fractions. Distinct tumor geometry of the subcomponents, irrespective of histologic grade, is potentially a phenotypic manifestation of the genetic and epigenetic landscape of the tumors. Indeed, WHO revised glioma classification to introduce a molecular basis of glioma classification as IDH mutant and wildtype, wherein an IDH mutant tumor of similar histologic grade presents with good prognosis, longer overall survival with better response to the treatment compared to the IDH wildtype gliomas. Similarly, a glioma harboring MGMT methylation is better suited for treatment with alkylating chemotherapy, while MGMT unmethylated gliomas are chemoresistant and are associated with poor survival. Determining IDH and MGMT status demands tumor tissue biopsy. Obtaining a tumor tissue biopsy via surgical interventions to determine molecular markers suggestive of the course of treatment and prognostication is one of the limiting factors for the improved clinical management of the patients. Additionally, it gets further complicated for the tumors that are in inaccessible locations like the brain stem. A noninvasive precise method of discriminating between IDH mutant *vs*. IDH wildtype tumors MGMT methylated *vs*. unmethylated tumors is needed for improved clinical management of glioma patients. Therefore, in this study, the geometric heterogeneity of the glioma subcomponents is quantified, in IDH mutant *vs*. IDH wildtype gliomas, and MGMT methylated *vs*. MGMT unmethylated, as FD and lacunarity using multimodal MR imaging data to establish a noninvasive quantitative radio-genomic platform depictive of IDH and MGMT status. Indeed, we find that FDs of enhancing and nonenhancing subcomponents are distinct in IDH mutant *vs*. wildtype, MGMT methylated, and unmethylated gliomas. While the tumor subcomponent fractality and lacunarity are definitive of IDH status, they are also associated with overall survival.

A significant fraction of enhancing subcomponents in glioma is a typical characteristic of aggressive and high-grade gliomas. Fractal dimensions of enhancing and nonenhancing subcomponents were distinct across mutational status. Our findings of increased fractality of the enhancing subcomponent and reduced fractality of the nonehnacing subcomponent in the IDH wildtype glioma group and its association with shortened overall survival (Figure 1C-H, Table 2A) indicates that the subcomponent geometry is potentially a manifestation of the molecular status and associated prognosis. Reduced fractality of the enhancing subcomponent, in an IDH mutant glioma is suggestive of smooth geometry for the enhancing subcomponent is a feature typical of IDH mutant tumors while highly irregular enhancing subcomponent appears to be a feature typical of IDH wildtype tumors. The higher fractality (irregular geometry) in the enhancing subcomponents associates with shortened overall survival; a feature typical of IDH wildtype gliomas. Irregular geometry of the enhancing subcomponents in IDH wildtype is likely to contribute to high structural heterogeneities; thereby explaining poor response to treatment in IDH wild type gliomas. Smooth enhancing subcomponent geometry and irregular nonenhancing geometry may potentially serve as a unique feature of IDH mutant tumors and longer overall survival. While our study did not reveal any significant disparities in the FD and lacunarity of the edema subcomponent between IDH mutant and wildtype gliomas, it is worth noting that the lacunarity of this particular component was still able to distinguish overall survival in the entire cohort.

Stratification of subjects for the MGMT epigenetic status across IDH mutant and wildtype clearly depicts that MGMT status does not have significant phenotypic impact on subcomponent geometry. IDH mutant gliomas with MGMT methylation does not differ for fractality or lacunarity when compared to IDH mutant MGMT unmethylated gliomas. Similarly, the IDH wildtype gliomas with MGMT methylation and unmethylations also did not differ for fractality and lacunarity. The difference was only noticed between IDH mutant and wildtype gliomas with either MGMT status. No differences in the fractality or lacunarity across the enhancing or nonenhancing subcomponents for MGMT status within a IDH type clearly depicts that fractality and lacunarity is a unique manifestation depictive of only IDH status.

When used as a single independent variable for IDH status classification, the FD of the subcomponents provided lower accuracies, wherein the edema subcomponent exhibits the lowest (∼50±6%). In contrast, the enhancing subcomponent exhibits the highest accuracy (∼80±8%) (Figure 3C). However, a combination of fractality of enhancing and nonenhancing subcomponents together serves as the most ‘optimal feature’ with the highest discriminative accuracy (∼87±8%) for IDH status across all three machine learning methods. The inclusion of the edema subcomponent does not alter the predictive accuracies of the models, reinforcing that IDH mutation status is predominantly associated with the geometry of enhancing and nonenhancing tumor subcomponents. While the combination of FD of enhancing and nonenhancing subcomponents has significantly poor accuracy in classifying tumors for MGMT status (Supplementary Figure S2 and S3), this establishes that distinct phenotypic manifestation of tumor geometry heterogeneity as fractality or lacunarity across tumors and its subcomponents will identify closely with a unique molecular status. Fractal dimension and lacunarity are independent geometric entities. While, in this study, they may appear to share an inverse relationship, they are not bound mathematically by any function of inverse relation. In certain fractal patterns, an increase in FD might correspond to a decrease in lacunarity, suggesting a more uniform space distribution. These metrics capture distinct aspects of complex systems; their association can be complex or unrelated.

Similarly, the combination of lacunarity of enhancing and nonenhancing subcomponents showed good accuracy in classifying IDH mutation status. This suggests the texture (irregularity) and progression pattern (patchiness) of tumor subcomponents, individually, are expressed as the observable characteristics expressed due to the underlying genetic constitute, which in this case is the IDH status.

Fractal dimension and lacunarity between IDH mutant and wildtype gliomas were not distinct, unlike the differences observed for enhancing and nonenhancing subcomponents. Edema is not a cellular entity but rather an accumulation of water because of displacement or death of normal cells while the tumor proliferates and grows; thereby, edema may not account for the phenotypic manifestation of any molecular subtype.

Studies in the past have attempted predicting the IDH status using tumor volumetry and pattern using ML models ^28,29^. Several investigations have examined glioma grade, progression, and prognostic implications in patients through volumetric analysis, as well as the assessment of textures and patterns within the entire tumor or specific components such as edema and nonenhancing regions (encompassing the necrotic core) ^17,30–33^. Overall volumetric measurements may not mimic the phenotypic manifestations, while the gliomas with similar volume will present with distinct shape and geometry. A volume measurement apparently is a cross product of surface area and thickness, therefore a subtle change in surface area and no change in thickness or vice versa may remain masked in the overall tumor volume estimates. Interestingly a volumetric study pertaining to aging associated changes suggested that surface area and thickness are governed have distinct genetic influences ^34^, henceforth measurement of tumor subcomponent geometry as fractal dimensions and lacunarity provide a unique quantitative basis of determining molecular subtypes of gliomas.

The stratified 5-fold cross-validation strategy was used to ensure that the machine learning models avoided any uncertainty arising from random sampling. For each fold of the prediction, the two sets were independent of each other and randomly selected from the entire cohort with the least possible skew of the dependent variables to avoid any artificial increase in the prediction accuracies. For a comprehensive assessment of the performance of the three models, confusion matrices were generated, in addition to ROC curves that provide a graphical representation of a model’s ability to discriminate between classes at various thresholds to provide a detailed interpretation of the model performance, taking into account the True Positive, True Negatives, False Positives, and False Negatives. The normalized confusion matrices of FD or lacunarity of the three subcomponents show increasing accuracy while distinguishing between the IDH wildtype and IDH mutant status when we consider each subcomponent individually and when the features of the subcomponents are combined as the independent variable for classification.

While this research demonstrated high accuracy in distinguishing IDH status across three distinct machine learning classifiers, the clinical context necessitates achieving 100 percent accuracy, given that treatment decisions hinge on the model’s predictions. To attain this objective, two fundamental steps are needed. First, expanding the dataset size is imperative, as deep learning algorithms excel with larger datasets, resulting in enhanced predictive capabilities. Second, comprehensive datasets encompassing tumor segmentation into four subcomponents based on tissue composition and data on molecular subtypes are essential for training models to effectively segment and classify gliomas.

The higher Fractal Dimension of the enhancing subcomponent presented with a poor prognosis when the whole cohort was divided based on the log rank statistic. The subset of subjects (∼92%) with an FD of the enhancing subcomponent > 0.69 and had shortened survival were the IDH wildtype, while ∼83% were IDH mutant in the cohort of FD < 0.69. The distinct survival based on cutoff further confirms the geometry associations with mutational status wherein lower FD of the enhancing subcomponent serves as a phenotypic manifestation of mutational status and shows a longer overall survival of 44 months. Similarly, for the nonenhancing subcomponent, the increased FD (>1.2) is associated with longer overall survival. Noticeably, around 82% of the subjects were IDH mutants for the subset of patients with FD > 1.2, which further explains that the survival benefit observed in subjects with higher FD for the nonenhancing subcomponent arises predominantly owing to the IDH mutation. Survival as a function of structural geometry is a phenotypic manifestation of genetic status as IDH mutant or wildtype. Higher FDs indicate increased irregularity, while lower FDs are associated with decreased irregularity. This study lays a quantitative platform that estimates tumor subcomponent geometry as a clinically precise, noninvasive, and easy-to-use benchmark of genotypic status and overall survival. In addition to understanding tumor structural heterogeneity and geometry as functions of genetic and epigenetic landscape, this study opens a plethora of questions regarding a need to understand tissue geometry and its functionality. A tissue may attain more irregularity for its advantage. In contrast, another tissue may become more homogeneous and smoother, which may be an underlying variable in understanding aging and other brain disorders.

## Authorship

V.T., N.Y., and V.A. conceptualized and designed the study; A.M., N.Y., and V.T. analyzed and interpreted the data, prepared the script (code) for analysis and figures; A.M., N.Y., and V.T. drafted the manuscript. N.M. helped initially in the literature search. All authors reviewed the results and approved the final version of the manuscript.

## Data Availability

All the Python and R codes and data used for statistical measurements can be provided upon request. In case of any clarifications, data can be directly provided upon request.

The imaging data was obtained from the TCIA portal (https://www.cancerimagingarchive.net/). The tumor subcomponent masks were obtained using the information published by Bakas S _et al._20,21

## Supporting information

Supplementary material all

## Acknowledgments

The authors acknowledge the support of TCIA for the imaging data and TCGA for the IDH and MGMT labels. The authors would also like to thank IISER Berhampur for the computational facility. The authors would also like to thanks Niraj Kumar Gupta for proofreading the article.

