## Supplementary material all for "Fractal Dimension and Lacunarity Measures of Glioma Subcomponents Provide a Quantitative Platform Discriminative of IDH Status: A Radiogenomics Approach in Gliomas"

Supplementary Table S1

| S. No. | Patient ID | Sex | Age at Diagnosis, y | Tumor Type† | Grade | IDH Status | MGMT Status# | Fractal Dimension (Mean) | | | Lacunarity (Mean) | | | Survival, *mo* |
|---|---|---|---|---|---|---|---|---|---|---|---|---|---|---|
| | | | | | | | | Enhancing | Nonenhancing | Edema | Enhancing | Nonenhancing | Edema | |
| 1 | TCGA-CS-4942 | F | 44 | A | G3 | Mutant | UM | 0.45 | 1.21 | 1.97 | 8.15 | 1.17 | 0.93 | 43.86 |
| 2 | TCGA-CS-4944 | M | 50 | A | G2 | Mutant | M | 0 | 1.37 | 1.97 | N/A* | 1.35 | 0.91 | 10.61 |
| 3 | TCGA-CS-5393 | M | 39 | A | G3 | Mutant | M | 0.49 | 1.23 | 2.00 | 8.20 | 1.24 | 0.81 | 40.15 |
| 4 | TCGA-CS-5396 | F | 53 | OG | G3 | Mutant | M | 1.79 | 1.79 | 1.42 | 1.86 | 1.54 | 21.47 | 9.96 |
| 5 | TCGA-CS-5397 | F | 54 | A | G3 | Wildtype | UM | 0 | 1.14 | 1.97 | N/A* | 1.52 | 1.03 | 6.37 |
| 6 | TCGA-CS-6186 | M | 58 | OA | G3 | Wildtype | UM | 0.77 | 0.93 | 1.95 | 5.01 | 4.50 | 0.96 | 17.68 |
| 7 | TCGA-CS-6188 | M | 48 | A | G3 | Wildtype | UM | 0.40 | 0.44 | 1.82 | 4.55 | 3.24 | 1.29 | 23.82 |
| 8 | TCGA-CS-6665 | F | 51 | A | G3 | Mutant | M | 0 | 1.30 | 1.91 | N/A* | 2.00 | 1.42 | 12.42 |
| 9 | TCGA-CS-6666 | M | 22 | A | G3 | Mutant | M | 0 | 1.48 | 1.87 | N/A* | 0.96 | 2.77 | 8.48 |
| 10 | TCGA-CS-6668 | F | 57 | OG | G2 | Mutant | M | 0.47 | 0.31 | 1.83 | 5.98 | 9.32 | 0.93 | 8.02 |
| 11 | TCGA-CS-6669 | F | 26 | OG | G2 | Wildtype | UM | 0 | 0.86 | 1.99 | N/A* | 1.79 | 0.80 | 7.39 |
| 12 | TCGA-DU-5851 | F | 40 | OA | G3 | Mutant | UM | 0.69 | 0.83 | 1.97 | 3.70 | 3.18 | 0.86 | 17.45 |
| 13 | TCGA-DU-5854 | F | 57 | A | G3 | Wildtype | UM | 1.18 | 0.21 | 1.95 | 2.05 | 6.02 | 0.93 | 8.44 |
| 14 | TCGA-DU-5855 | F | 49 | OA | G3 | Mutant | M | 0.97 | 1.65 | 1.95 | 8.51 | 0.97 | 1.25 | 6.80 |
| 15 | TCGA-DU-5872 | F | 43 | OA | G2 | Mutant | M | 1.16 | 1.56 | 1.99 | 5.22 | 0.93 | 1.42 | 17.48 |
| 16 | TCGA-DU-5874 | F | 62 | OG | G2 | Mutant | M | 0 | 1.35 | 1.72 | N/A* | 0.69 | 2.71 | 15.15 |
| 17 | TCGA-DU-6404 | F | 24 | OG | G3 | Wildtype | UM | 1.52 | 0.66 | 1.91 | 0.93 | 2.81 | 0.98 | 133.65 |
| 18 | TCGA-DU-6542 | M | 25 | OA | G3 | Mutant | M | 1.39 | 1.43 | 1.89 | 2.11 | 1.46 | 1.16 | 7.72 |
| 19 | TCGA-DU-7008 | F | 41 | OG | G2 | Mutant | M | 0 | 1.59 | 2.07 | N/A* | 2.76 | 0.88 | 156.13 |
| 20 | TCGA-DU-7010 | F | 58 | A | G3 | Mutant | M | 0.53 | 0.65 | 1.97 | 7.99 | 6.57 | 0.79 | 14.98 |
| 21 | TCGA-DU-7015 | F | 41 | OG | G2 | Mutant | M | 0.96 | 1.50 | 1.93 | 3.53 | 1.10 | 1.23 | 90.71 |
| 22 | TCGA-DU-7018 | F | 57 | OG | G3 | Mutant | M | 1.44 | 1.59 | 1.97 | 4.07 | 1.57 | 1.73 | 30.65 |
| 23 | TCGA-DU-7019 | M | 39 | OA | G3 | Mutant | M | 0 | 1.40 | 1.97 | N/A* | 1.41 | 1.28 | 26.28 |
| 24 | TCGA-DU-7294 | F | 53 | OG | G2 | Mutant | M | 1.10 | 1.43 | 1.94 | 5.99 | 1.30 | 1.01 | 94.26 |
| 25 | TCGA-DU-7298 | F | 38 | A | G3 | Mutant | M | 1.26 | 1.56 | 2.03 | 3.52 | 1.03 | 0.96 | 18.92 |
| 26 | TCGA-DU-7299 | M | 33 | A | G3 | Mutant | M | 0 | 1.07 | 2.03 | N/A* | 2.37 | 1.38 | 43.99 |
| 27 | TCGA-DU-7300 | F | 53 | OG | G3 | Mutant | M | 0.37 | 0.98 | 1.93 | 9.60 | 1.18 | 0.75 | 61.96 |
| 28 | TCGA-DU-7301 | M | 53 | OG | G2 | Mutant | M | 0.50 | 1.53 | 1.94 | 10.48 | 0.81 | 1.58 | 25.89 |

| | | | | | | | | | | | | | | |
|---|---|---|---|---|---|---|---|---|---|---|---|---|---|---|
| 29 | TCGA-DU-7302 | F | 48 | OG | G3 | Mutant | M | 0.51 | 1.30 | 1.85 | 5.82 | 1.33 | 1.38 | 60.26 |
| 30 | TCGA-DU-7304 | M | 43 | OA | G3 | Mutant | M | 0.79 | 1.01 | 1.98 | 4.50 | 1.14 | 0.86 | 23.29 |
| 31 | TCGA-DU-7306 | M | 67 | OA | G2 | Mutant | M | 0.80 | 1.83 | 1.51 | 5.45 | 0.62 | 3.10 | 41.96 |
| 32 | TCGA-DU-7309 | F | 41 | OG | G3 | Mutant | M | 0.54 | 1.93 | 1.37 | 2.88 | 1.17 | 1.89 | 2.76 |
| 33 | TCGA-DU-8162 | F | 61 | OA | G3 | Wildtype | UM | 0 | 1.71 | 1.44 | N/A* | 0.99 | 3.71 | 14.59 |
| 34 | TCGA-DU-8164 | M | 51 | OG | G2 | Mutant | M | 0 | 1.52 | 1.92 | N/A* | 0.87 | 0.94 | 21.39 |
| 35 | TCGA-DU-8166 | F | 29 | OA | G2 | Mutant | M | 0 | 1.72 | 1.90 | N/A* | 0.95 | 1.22 | 16.95 |
| 36 | TCGA-DU-8167 | F | 69 | OA | G2 | Mutant | M | 0.08 | 1.68 | 1.96 | 10.54 | 0.97 | 1.47 | 15.47 |
| 37 | TCGA-DU-8168 | F | 55 | OG | G3 | Mutant | M | 0.94 | 1.68 | 1.76 | 5.84 | 1.59 | 3.58 | 14.16 |
| 38 | TCGA-DU-A5TR | M | 51 | OA | G2 | Mutant | M | 0.89 | 1.70 | 1.86 | 8.24 | 0.74 | 3.90 | 12.58 |
| 39 | TCGA-DU-A5TS | M | 42 | OG | G2 | Mutant | M | 0 | 1.47 | 2.05 | N/A* | 1.16 | 1.06 | 15.11 |
| 40 | TCGA-DU-A5TT | M | 70 | OG | G3 | Wildtype | M | 0.42 | 1.38 | 1.97 | 8.34 | 2.52 | 1.17 | 4.96 |
| 41 | TCGA-DU-A5TU | F | 62 | A | G2 | Mutant | M | 0 | 1.55 | 1.92 | N/A* | 1.26 | 0.89 | 3.65 |
| 42 | TCGA-DU-A5TW | F | 33 | A | G3 | Mutant | M | 0.40 | 1.53 | 1.99 | 7.95 | 1.35 | 0.83 | 5.65 |
| 43 | TCGA-DU-A6S7 | F | 27 | A | G3 | Mutant | M | 0.19 | 1.54 | 1.98 | 7.08 | 1.15 | 1.46 | 7.23 |
| 44 | TCGA-DU-A6S8 | F | 74 | OG | G3 | Mutant | M | 0.55 | 1.41 | 1.88 | 12.38 | 2.38 | 1.42 | 5.98 |
| 45 | TCGA-FG-5964 | M | 62 | OG | G2 | Mutant | M | 1.10 | 1.16 | 1.89 | 3.05 | 2.41 | 1.17 | 34.33 |
| 46 | TCGA-FG-6689 | M | 30 | A | G2 | Mutant | M | 0.57 | 1.74 | 1.96 | 11.59 | 1.19 | 1.74 | 14.92 |
| 47 | TCGA-FG-6691 | F | 23 | A | G2 | Mutant | UM | 0 | 1.44 | 1.96 | N/A* | 0.92 | 1.25 | 23.95 |
| 48 | TCGA-FG-6692 | M | 63 | OG | G3 | Wildtype | M | 0.92 | 0.68 | 1.85 | 1.85 | 3.47 | 0.84 | 18.43 |
| 49 | TCGA-FG-7634 | M | 28 | OG | G2 | Mutant | M | 0 | 1.27 | 1.97 | N/A* | 0.88 | 0.72 | 15.34 |
| 50 | TCGA-HT-7473 | M | 28 | OA | G2 | Mutant | UM | 0.53 | 1.28 | 1.93 | 6.20 | 4.07 | 0.89 | 16.53 |
| 51 | TCGA-HT-7602 | M | 21 | OG | G2 | Mutant | M | 0 | 1.48 | 1.80 | N/A* | 0.98 | 2.10 | 29.83 |
| 52 | TCGA-HT-7680 | F | 32 | A | G2 | Wildtype | UM | 0.26 | 1.48 | 1.97 | 7.32 | 2.00 | 1.74 | 0.76 |
| 53 | TCGA-HT-7686 | F | 29 | A | G3 | Mutant | M | 0.66 | 1.85 | 0 | 7.53 | 0.73 | N/A* | 42.71 |
| 54 | TCGA-HT-7690 | M | 29 | OA | G3 | Mutant | M | 1.05 | 1.88 | 1.87 | 3.44 | 0.82 | 2.44 | 0.10 |
| 55 | TCGA-HT-7694 | M | 60 | OG | G3 | Mutant | M | 0.19 | 1.42 | 1.97 | 11.97 | 1.33 | 1.36 | 6.90 |
| 56 | TCGA-HT-7879 | M | 31 | OA | G3 | Mutant | M | 0 | 1.45 | 1.89 | N/A* | 0.98 | 1.98 | 3.68 |
| 57 | TCGA-HT-7884 | F | 44 | A | G2 | Mutant | M | 0 | 1.60 | 1.95 | N/A* | 1.20 | 1.42 | 11.27 |
| 58 | TCGA-HT-8018 | F | 40 | OA | G2 | Mutant | M | 0.53 | 0.85 | 1.90 | 2.76 | 1.26 | 0.82 | 21.49 |
| 59 | TCGA-HT-8114 | M | 36 | OA | G3 | Mutant | M | 1.19 | 1.81 | 1.60 | 4.77 | 0.81 | 4.36 | 3.88 |
| 60 | TCGA-HT-8563 | F | 30 | A | G3 | Mutant | UM | 1.01 | 0.64 | 1.94 | 3.16 | 4.25 | 0.79 | 16.03 |
| 61 | TCGA-HT-A61A | F | 20 | OG | G2 | Mutant | M | 0 | 1.41 | 1.91 | N/A* | 1.47 | 1.13 | 6.37 |
| 62 | TCGA-02-0006 | F | 56 | GBM | G4 | Wildtype | UM | 0.93 | 0.61 | 1.96 | 7.91 | 7.86 | 0.94 | 18.33 |
| 63 | TCGA-02-0009 | F | 61 | GBM | G4 | Wildtype | UM | 0.92 | 1.01 | 1.89 | 2.52 | 2.39 | 0.86 | 10.58 |
| 64 | TCGA-02-0011 | F | 18 | GBM | G4 | Wildtype | M | 1.44 | 1.49 | 1.78 | 1.72 | 1.29 | 1.34 | 20.70 |
| 65 | TCGA-02-0027 | F | 33 | GBM | G4 | Wildtype | UM | 1.33 | 1.12 | 1.98 | 1.59 | 2.35 | 0.93 | 12.16 |

| | | | | | | | | | | | | | | |
|---|---|---|---|---|---|---|---|---|---|---|---|---|---|---|
| 66 | TCGA-02-0033 | M | 54 | GBM | G4 | Wildtype | M | 1.31 | 0.76 | 1.92 | 2.24 | 5.65 | 1.44 | 2.83 |
| 67 | TCGA-02-0034 | M | 60 | GBM | G4 | Wildtype | UM | 1.42 | 1.19 | 1.92 | 1.34 | 1.79 | 0.83 | 14.13 |
| 68 | TCGA-02-0037 | F | 74 | GBM | G4 | Wildtype | UM | 1.57 | 1.01 | 2.05 | 1.28 | 3.03 | 0.89 | 3.61 |
| 69 | TCGA-02-0046 | M | 61 | GBM | G4 | Wildtype | M | 1.04 | 0.84 | 1.91 | 1.82 | 3.29 | 0.97 | 6.87 |
| 70 | TCGA-02-0047 | M | 78 | GBM | G4 | Wildtype | UM | 1.05 | 0.75 | 2.00 | 2.36 | 4.30 | 0.85 | 14.72 |
| 71 | TCGA-02-0054 | F | 44 | GBM | G4 | Wildtype | M | 1.11 | 1.19 | 1.97 | 3.28 | 2.13 | 0.95 | 6.54 |
| 72 | TCGA-02-0064 | M | 50 | GBM | G4 | Wildtype | M | 1.76 | 1.23 | 1.85 | 1.20 | 2.18 | 1.65 | 19.71 |
| 73 | TCGA-02-0068 | M | 57 | GBM | G4 | Wildtype | N/A* | 1.18 | 1.19 | 1.91 | 1.53 | 2.01 | 0.76 | 26.42 |
| 74 | TCGA-02-0069 | F | 31 | GBM | G4 | Wildtype | M | 1.45 | 1.43 | 1.99 | 2.28 | 1.47 | 1.07 | 28.68 |
| 75 | TCGA-02-0075 | M | 63 | GBM | G4 | Wildtype | M | 1.10 | 1.14 | 1.78 | 2.31 | 1.21 | 1.91 | 20.83 |
| 76 | TCGA-02-0085 | F | 63 | GBM | G4 | Wildtype | M | 1.55 | 0.84 | 1.91 | 3.37 | 4.63 | 0.90 | 51.29 |
| 77 | TCGA-02-0086 | F | 45 | GBM | G4 | Wildtype | UM | 1.52 | 1.17 | 2.01 | 2.39 | 5.15 | 1.94 | 8.81 |
| 78 | TCGA-02-0102 | M | 42 | GBM | G4 | Wildtype | UM | 0.61 | 0.55 | 1.92 | 3.09 | 4.28 | 0.83 | 27.01 |
| 79 | TCGA-02-0116 | M | 51 | GBM | G4 | Wildtype | UM | 1.50 | 0.73 | 1.82 | 0.84 | 3.31 | 0.93 | 48.92 |
| 80 | TCGA-06-0119 | F | 81 | GBM | G4 | Wildtype | M | 1.17 | 0.82 | 1.93 | 0.81 | 2.15 | 1.04 | 2.69 |
| 81 | TCGA-06-0122 | F | 84 | GBM | G4 | Wildtype | UM | 0.97 | 0.67 | 1.87 | 3.47 | 5.26 | 1.31 | 6.14 |
| 82 | TCGA-06-0130 | M | 54 | GBM | G4 | Wildtype | UM | 1.11 | 0.78 | 1.84 | 1.91 | 3.70 | 0.95 | 12.94 |
| 83 | TCGA-06-0137 | F | 63 | GBM | G4 | Wildtype | UM | 1.41 | 1.16 | 1.83 | 1.30 | 3.38 | 0.99 | 26.68 |
| 84 | TCGA-06-0138 | M | 43 | GBM | G4 | Wildtype | N/A* | 1.43 | 1.27 | 1.78 | 1.61 | 2.28 | 1.99 | 24.21 |
| 85 | TCGA-06-0139 | M | 40 | GBM | G4 | Wildtype | UM | 1.58 | 1.53 | 1.91 | 1.56 | 2.07 | 1.56 | 11.89 |
| 86 | TCGA-06-0142 | M | 81 | GBM | G4 | Wildtype | UM | 1.51 | 1.42 | 1.94 | 2.93 | 2.45 | 0.89 | 2.20 |
| 87 | TCGA-06-0145 | F | 53 | GBM | G4 | Wildtype | M | 1.61 | 1.45 | 1.87 | 1.75 | 1.66 | 1.28 | 2.33 |
| 88 | TCGA-06-0154 | M | 54 | GBM | G4 | Wildtype | N/A* | 1.44 | 1.34 | 1.83 | 1.43 | 2.41 | 0.82 | 13.93 |
| 89 | TCGA-06-0158 | M | 73 | GBM | G4 | Wildtype | N/A* | 1.10 | 0.58 | 1.97 | 2.13 | 4.62 | 0.91 | 10.81 |
| 90 | TCGA-06-0176 | M | 34 | GBM | G4 | Wildtype | N/A* | 1.36 | 1.26 | 1.94 | 1.66 | 2.21 | 1.07 | 51.32 |
| 91 | TCGA-06-0184 | M | 63 | GBM | G4 | Wildtype | N/A* | 1.40 | 1.23 | 1.89 | 1.50 | 2.48 | 1.21 | 40.35 |
| 92 | TCGA-06-0185 | M | 54 | GBM | G4 | Wildtype | N/A* | 1.71 | 1.72 | 1.85 | 4.66 | 6.14 | 1.07 | 36.99 |
| 93 | TCGA-06-0187 | M | 69 | GBM | G4 | Wildtype | N/A* | 1.54 | 1.17 | 1.96 | 2.65 | 3.71 | 1.09 | 27.20 |
| 94 | TCGA-06-0188 | M | 71 | GBM | G4 | Wildtype | N/A* | 0.93 | 0.20 | 1.93 | 4.35 | 9.81 | 1.02 | 28.45 |
| 95 | TCGA-06-0190 | M | 62 | GBM | G4 | Wildtype | N/A* | 0.97 | 0.83 | 2.02 | 2.54 | 5.05 | 0.97 | 10.42 |
| 96 | TCGA-06-0192 | M | 58 | GBM | G4 | Wildtype | N/A* | 1.37 | 1.34 | 1.94 | 1.85 | 2.74 | 1.50 | 18.30 |
| 97 | TCGA-06-0240 | M | 57 | GBM | G4 | Wildtype | N/A* | 1.27 | 0.57 | 1.96 | 1.23 | 3.45 | 1.08 | 20.40 |
| 98 | TCGA-06-0644 | M | 71 | GBM | G4 | Wildtype | N/A* | 0.81 | 0.58 | 1.88 | 5.03 | 8.39 | 1.46 | 12.32 |
| 99 | TCGA-06-0646 | M | 60 | GBM | G4 | Wildtype | N/A* | 0.79 | 0.43 | 1.88 | 2.32 | 3.81 | 0.86 | 5.75 |
| 100 | TCGA-06-1084 | M | 54 | GBM | G4 | Wildtype | M | 1.20 | 0.85 | 1.87 | 1.96 | 4.29 | 1.04 | 7.43 |
| 101 | TCGA-06-2570 | F | 21 | GBM | G4 | Mutant | M | 1.80 | 1.54 | 1.95 | 1.63 | 1.38 | 1.69 | 9.36 |
| 102 | TCGA-06-5408 | F | 54 | GBM | G4 | Wildtype | UM | 1.24 | 1.04 | 1.94 | 2.03 | 3.56 | 1.19 | 11.73 |

| | | | | | | | | | | | | | | |
|---|---|---|---|---|---|---|---|---|---|---|---|---|---|---|
| 103 | TCGA-06-5413 | M | 67 | GBM | G4 | Wildtype | UM | 1.23 | 1.07 | 1.97 | 2.26 | 2.80 | 1.16 | 8.81 |
| 104 | TCGA-06-5417 | F | 45 | GBM | G4 | Mutant | M | 1.91 | 1.88 | 1.87 | 1.92 | 1.30 | 1.35 | 5.09 |
| 105 | TCGA-06-6389 | F | 49 | GBM | G4 | Mutant | M | 1.02 | 0.99 | 1.83 | 1.66 | 1.62 | 0.70 | 7.79 |
| 106 | TCGA-08-0355 | F | 30 | GBM | G4 | Wildtype | N/A* | 1.41 | 1.00 | 1.68 | 2.10 | 2.78 | 7.08 | 24.54 |
| 107 | TCGA-08-0356 | F | 59 | GBM | G4 | Wildtype | N/A* | 1.37 | 1.09 | 1.85 | 1.04 | 1.95 | 1.32 | 31.08 |
| 108 | TCGA-08-0359 | F | 59 | GBM | G4 | Wildtype | N/A* | 1.72 | 1.55 | 2.00 | 1.52 | 1.30 | 1.16 | 3.38 |
| 109 | TCGA-08-0360 | M | 76 | GBM | G4 | Wildtype | N/A* | 1.57 | 1.30 | 1.94 | 2.61 | 2.21 | 1.53 | 15.38 |
| 110 | TCGA-08-0385 | M | 71 | GBM | G4 | Wildtype | N/A* | 1.81 | 1.48 | 1.87 | 1.09 | 1.49 | 2.07 | 2.69 |
| 111 | TCGA-08-0389 | M | 59 | GBM | G4 | Wildtype | N/A* | 1.14 | 0.92 | 1.96 | 1.46 | 2.14 | 0.64 | 15.34 |
| 112 | TCGA-08-0390 | M | 69 | GBM | G4 | Wildtype | N/A* | 1.04 | 0.69 | 1.93 | 4.00 | 2.38 | 0.69 | 13.96 |
| 113 | TCGA-08-0392 | M | 60 | GBM | G4 | Wildtype | N/A* | 1.79 | 1.53 | 1.82 | 1.82 | 2.68 | 1.86 | 0.72 |
| 114 | TCGA-12-0616 | F | 36 | GBM | G4 | Wildtype | N/A* | 1.86 | 1.62 | 1.93 | 1.09 | 1.85 | 1.83 | 14.72 |
| 115 | TCGA-12-0829 | M | 75 | GBM | G4 | Wildtype | M | 0.98 | 0.86 | 1.89 | 2.56 | 6.34 | 1.09 | 20.57 |
| 116 | TCGA-12-1598 | F | 75 | GBM | G4 | Wildtype | M | 1.21 | 0.98 | 1.88 | 1.60 | 3.41 | 0.73 | 15.64 |
| 117 | TCGA-12-1601 | M | 71 | GBM | G4 | Wildtype | UM | 1.18 | 0.98 | 1.97 | 4.55 | 2.92 | 1.11 | N/A* |
| 118 | TCGA-14-1456 | M | 23 | GBM | G4 | Mutant | UM | 1.31 | 1.60 | 2.01 | 5.10 | 1.48 | 1.65 | 40.94 |
| 119 | TCGA-14-1794 | M | 59 | GBM | G4 | Wildtype | UM | 1.86 | 1.47 | 1.92 | 2.20 | 1.81 | 2.72 | 0.99 |
| 120 | TCGA-14-1825 | M | 70 | GBM | G4 | Wildtype | UM | 1.42 | 1.13 | 1.96 | 1.61 | 2.68 | 1.14 | 7.62 |
| 121 | TCGA-14-3477 | F | 38 | GBM | G4 | Wildtype | UM | 1.07 | 1.13 | 1.94 | 2.91 | 3.07 | 0.77 | 3.78 |
| 122 | TCGA-19-1789 | F | 69 | GBM | G4 | Wildtype | M | 1.08 | 1.09 | 1.92 | 3.08 | 2.20 | 0.95 | 3.25 |
| 123 | TCGA-19-2624 | M | 51 | GBM | G4 | Wildtype | UM | 1.12 | 1.24 | 1.92 | 3.72 | 2.07 | 0.75 | 0.16 |
| 124 | TCGA-19-2631 | F | 74 | GBM | G4 | Wildtype | M | 1.61 | 1.60 | 1.95 | 2.48 | 5.76 | 0.68 | 7.00 |
| 125 | TCGA-19-5951 | F | 76 | GBM | G4 | Wildtype | UM | 1.72 | 1.58 | 1.81 | 1.03 | 2.39 | 1.54 | 8.02 |
| 126 | TCGA-19-5954 | F | 72 | GBM | G4 | Wildtype | M | 0.83 | 0.61 | 1.80 | 1.68 | 5.38 | 0.69 | 7.95 |
| 127 | TCGA-19-5958 | M | 56 | GBM | G4 | Wildtype | UM | 1.22 | 1.13 | 1.95 | 1.41 | 1.47 | 0.84 | 5.39 |
| 128 | TCGA-19-5960 | M | 56 | GBM | G4 | Wildtype | UM | 1.51 | 1.30 | 1.97 | 0.92 | 3.55 | 0.93 | 5.42 |
| 129 | TCGA-76-4932 | F | 50 | GBM | G4 | Wildtype | M | 1.58 | 1.62 | 1.94 | 1.76 | 1.67 | 2.04 | 47.90 |
| 130 | TCGA-76-4934 | F | 66 | GBM | G4 | Wildtype | M | 1.49 | 1.36 | 1.97 | 1.39 | 2.39 | 1.36 | 2.53 |
| 131 | TCGA-76-4935 | F | 52 | GBM | G4 | Wildtype | M | 1.84 | 1.40 | 1.92 | 1.22 | 1.37 | 1.65 | 10.78 |
| 132 | TCGA-76-6191 | M | 57 | GBM | G4 | Wildtype | UM | 1.41 | 1.12 | 1.92 | 1.44 | 2.50 | 0.82 | 16.69 |
| 133 | TCGA-76-6193 | M | 78 | GBM | G4 | Wildtype | UM | 0.76 | 0.10 | 1.93 | 4.81 | 15.42 | 0.80 | 2.69 |
| 134 | TCGA-76-6280 | M | 57 | GBM | G4 | Wildtype | M | 1.21 | 1.08 | 1.87 | 1.49 | 2.91 | 0.88 | 11.37 |
| 135 | TCGA-76-6282 | M | 63 | GBM | G4 | Wildtype | UM | 1.28 | 1.00 | 1.97 | 2.46 | 2.55 | 1.20 | 17.05 |
| 136 | TCGA-76-6285 | F | 64 | GBM | G4 | Wildtype | UM | 1.54 | 1.62 | 1.95 | 2.31 | 1.99 | 1.48 | 8.35 |
| 137 | TCGA-76-6656 | M | 66 | GBM | G4 | Wildtype | M | 1.54 | 1.27 | 1.95 | 1.76 | 3.35 | 0.81 | 4.83 |
| 138 | TCGA-76-6657 | M | 74 | GBM | G4 | Wildtype | M | 1.47 | 1.34 | 1.95 | 1.24 | 2.15 | 1.22 | 5.03 |
| 139 | TCGA-76-6661 | M | 54 | GBM | G4 | Wildtype | UM | 1.50 | 1.47 | 1.84 | 1.06 | 1.47 | 1.24 | 0.23 |

| | | | | | | | | | | | | | | |
|---|---|---|---|---|---|---|---|---|---|---|---|---|---|---|
| 140 | TCGA-76-6662 | M | 58 | GBM | G4 | Wildtype | UM | 1.00 | 0.79 | 1.89 | 2.84 | 3.25 | 1.22 | 9.27 |
| 141 | TCGA-76-6663 | F | 44 | GBM | G4 | Wildtype | UM | 1.37 | 1.38 | 1.99 | 2.91 | 2.00 | 1.66 | 7.72 |
| 142 | TCGA-76-6664 | F | 49 | GBM | G4 | Wildtype | M | 1.05 | 0.96 | 1.90 | 3.88 | 4.49 | 0.99 | 7.79 |

**† A: Astrocytoma, OA: Oligoastrocytoma, OG: Oligodendroglioma, GBM: Glioblastoma Multiforme**

*** Not available / Not calculated**

**# UM: Unmethylated, M: Methylated**

**Supplementary Figure S1.**

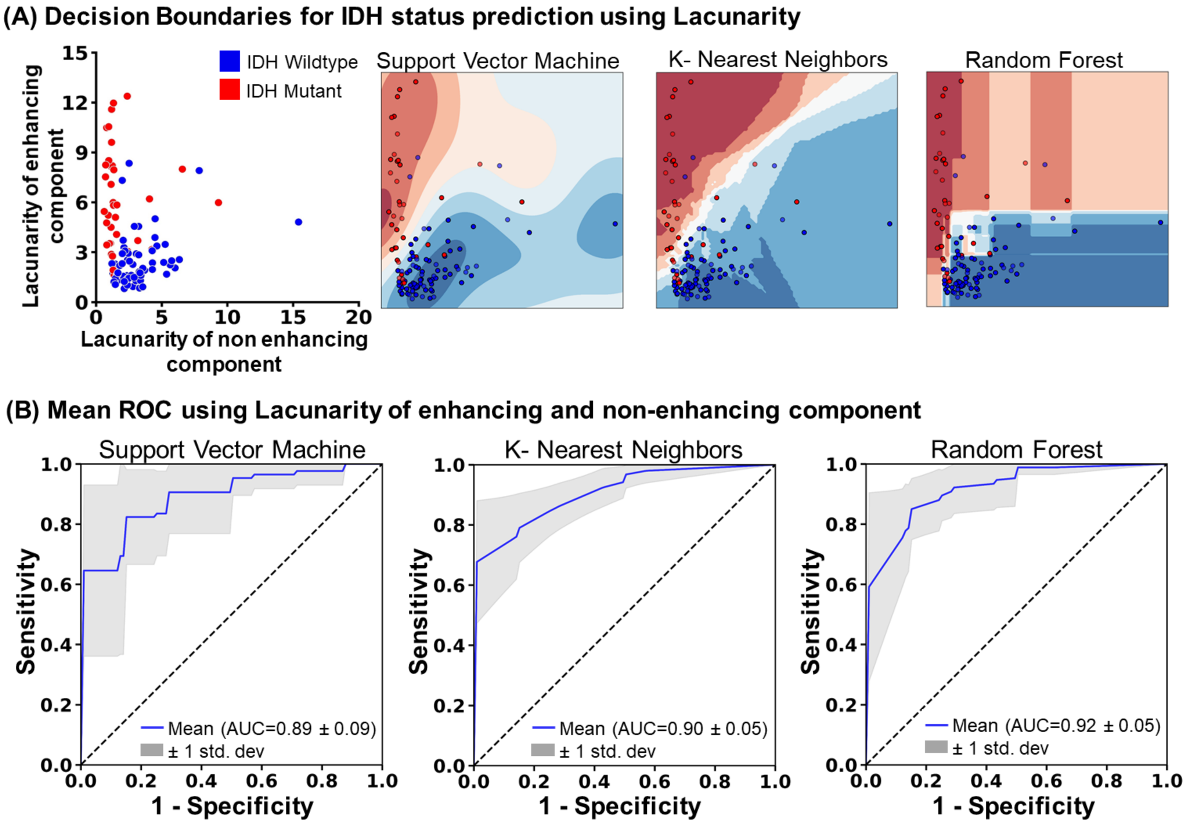


**Supplementary Figure S1: Machine learning models for differentiating IDH status using lacunarity**: (A) Scatter plot depicting the distribution of IDH mutation status (Red - Mutant, Blue - Wildtype) lacunarity of enhancing and nonenhancing tumor subcomponents. Decision boundaries of three machine learning algorithms- SVM, RF, and KNN discriminating the IDH status. (B) The Receiver Operating Characteristic (ROC) curve of the three ML- models: SVM, RF, and KNN, with mean AUC using the lacunarity of enhancing and nonenhancing tumor subcomponents for distinguishing IDH molecular status.

**Supplementary Figure S2.**

**(A) Decision Boundaries for MGMT status prediction using Fractal Dimension**

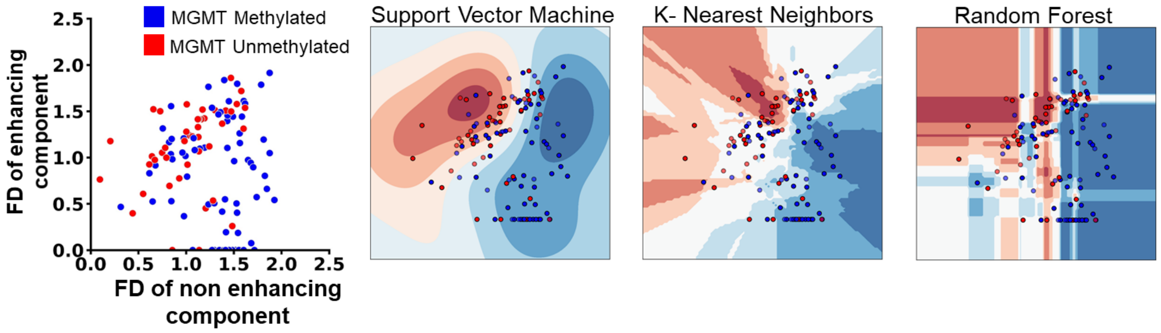


**(B) Mean ROC using Fractal Dimension of enhancing and non-enhancing component**

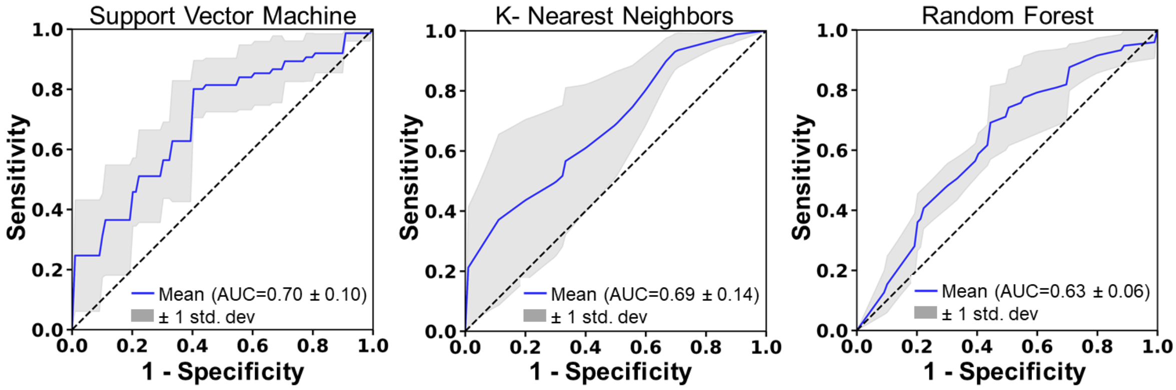


**Supplementary Figure S2: Machine learning models for differentiating MGMT status using fractal dimension.** (A) Scatter plot depicting the distribution of MGMT methylation status (Red -Unmethylated, Blue - Methylated) FD of enhancing and nonenhancing tumor subcomponents. Decision boundaries of three machine learning algorithms- SVM, RF, and KNN discriminating the MGMT status. (B) The Receiver Operating Characteristic (ROC) curve of the three ML- models: SVM, RF, and KNN, with mean AUC using the Fractal Dimension of enhancing and nonenhancing tumor subcomponents for distinguishing MGMT status.

**Supplementary Figure S3.**

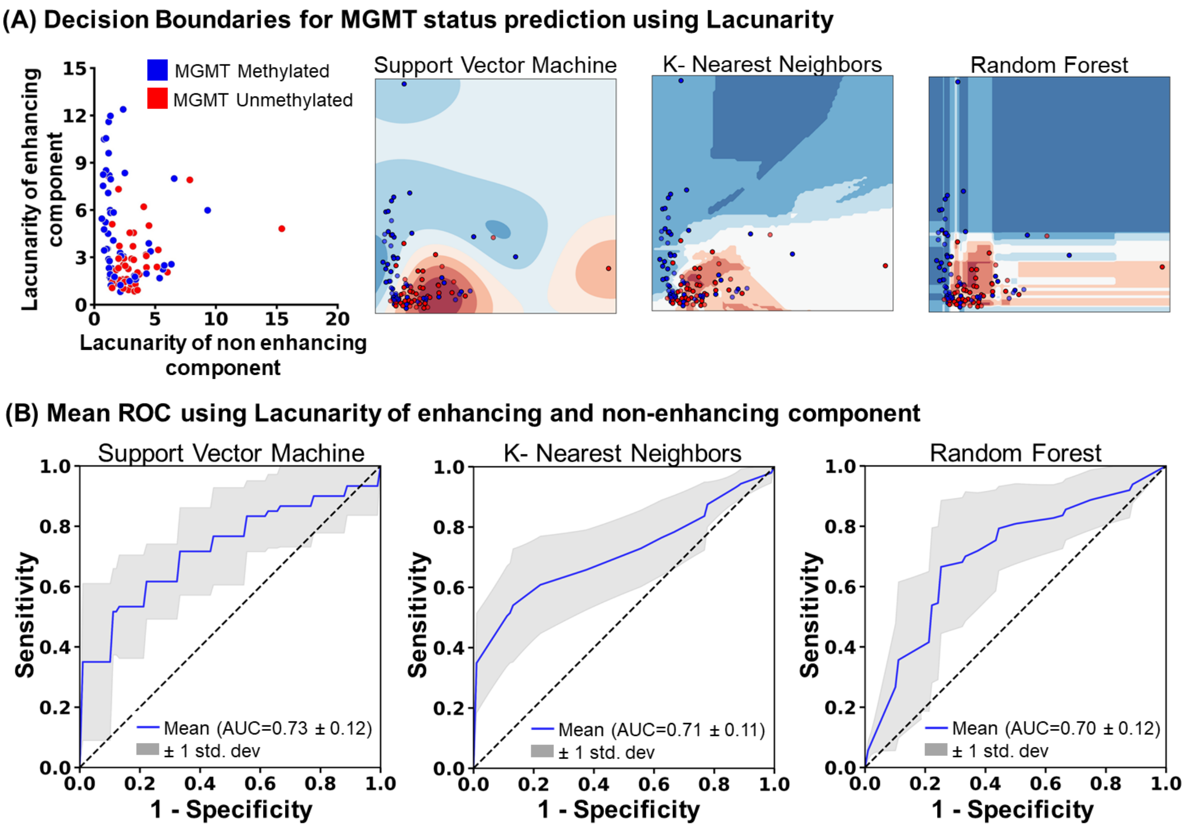


**Supplementary Figure S3: Machine learning models for differentiating MGMT status using lacunarity.** (A) Scatter plot depicting the distribution of MGMT methylation status (Red -Unmethylated, Blue - Methylated) lacunarity of enhancing and nonenhancing tumor subcomponents. Decision boundaries of three machine learning algorithms- SVM, RF, and KNN discriminating the MGMT status. (B) The Receiver Operating Characteristic (ROC) curve of the three ML-models: SVM, RF, and KNN, with mean AUC using the lacunarity of enhancing and nonenhancing tumor subcomponents for distinguishing MGMT status.

**Supplementary Figure S4.**

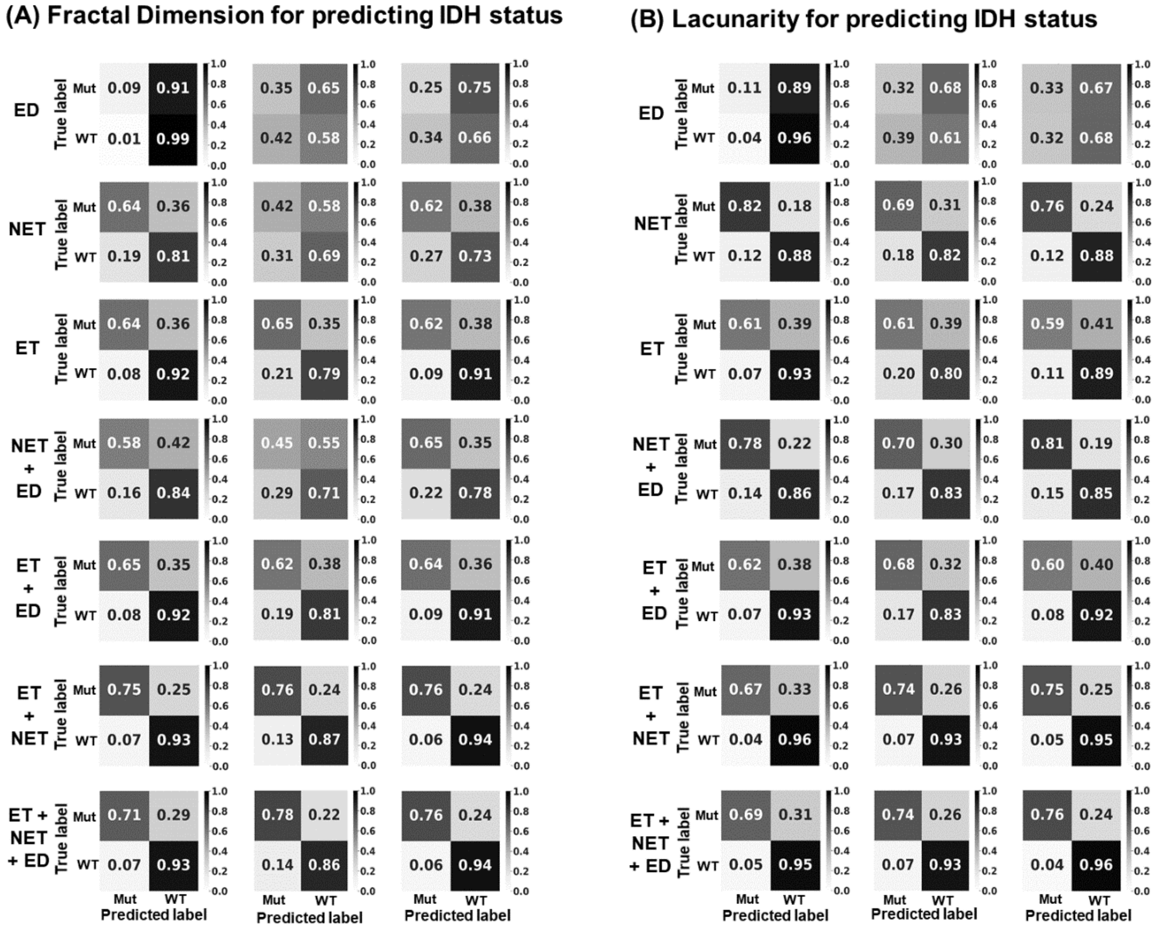


**Supplementary Figure S4: Normalized confusion matrices for IDH.** Normalized confusion matrices generated by the machine learning models (SVM, KNN, and RF) while classifying IDH status using the (A) Fractal dimension and (B) Lacunarity of each tumor subcomponent individually and in combination.

ET: Enhancing tumor region, NET: Nonenhancing tumor region, ED: Edema tumor region, Mut: Mutant (IDH), WT: wildtype (IDH)

**Supplementary Figure S5.**

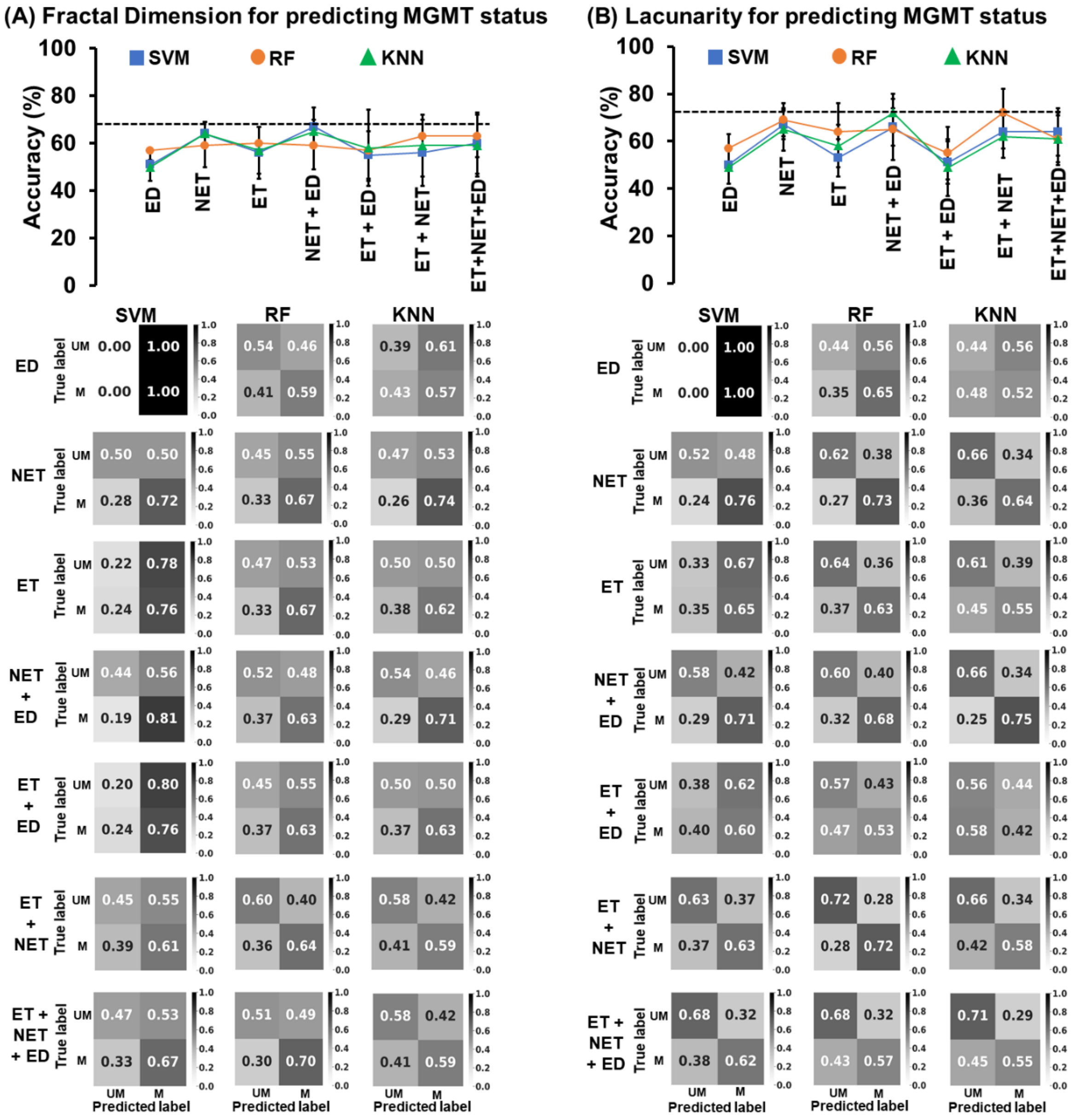


**Supplementary Figure S5: Accuracy plot and normalized confusion matrices for MGMT**. Accuracy plots accompanied by the normalized confusion matrices generated by the machine learning models (SVM, KNN, and RF) while classifying MGMT status using the (A) Fractal dimension and (B) Lacunarity of each tumor subcomponent individually and in combination. ET: Enhancing tumor region, NET: Nonenhancing tumor region, ED: Edema tumor region, UM: Unmethylated (MGMT), M: Methylated (MGMT)

**Supplementary Figure S6.**

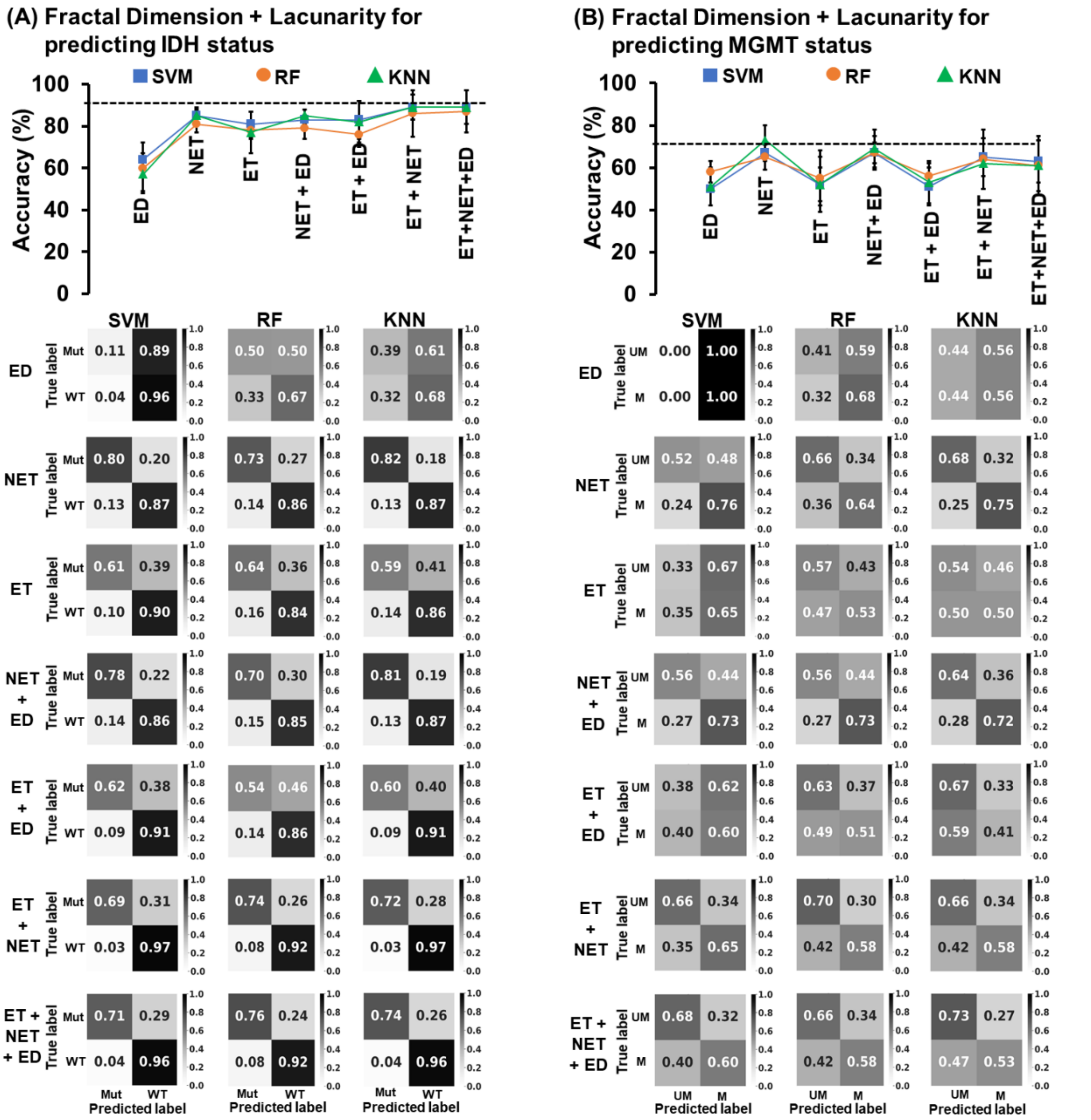


**Supplementary Figure S6: Accuracy plot and normalized confusion matrices for IDH and MGMT**. Accuracy plots accompanied by the normalized confusion matrices generated by the machine learning models (SVM, KNN, and RF) while classifying (A) IDH and (B) MGMT status using the Fractal dimension and Lacunarity (in combination) of each tumor subcomponent individually and combined. The width of the violin plot shows the approximate frequency of data points in that region shows the kernel density depictive of probability of finding a data.

ET: Enhancing tumor region, NET: Nonenhancing tumor region, ED: Edema tumor region, Mut: Mutant (IDH), WT: wildtype (IDH), UM: Unmethylated (MGMT), M: Methylated (MGMT)

**Supplementary Figure S7.**

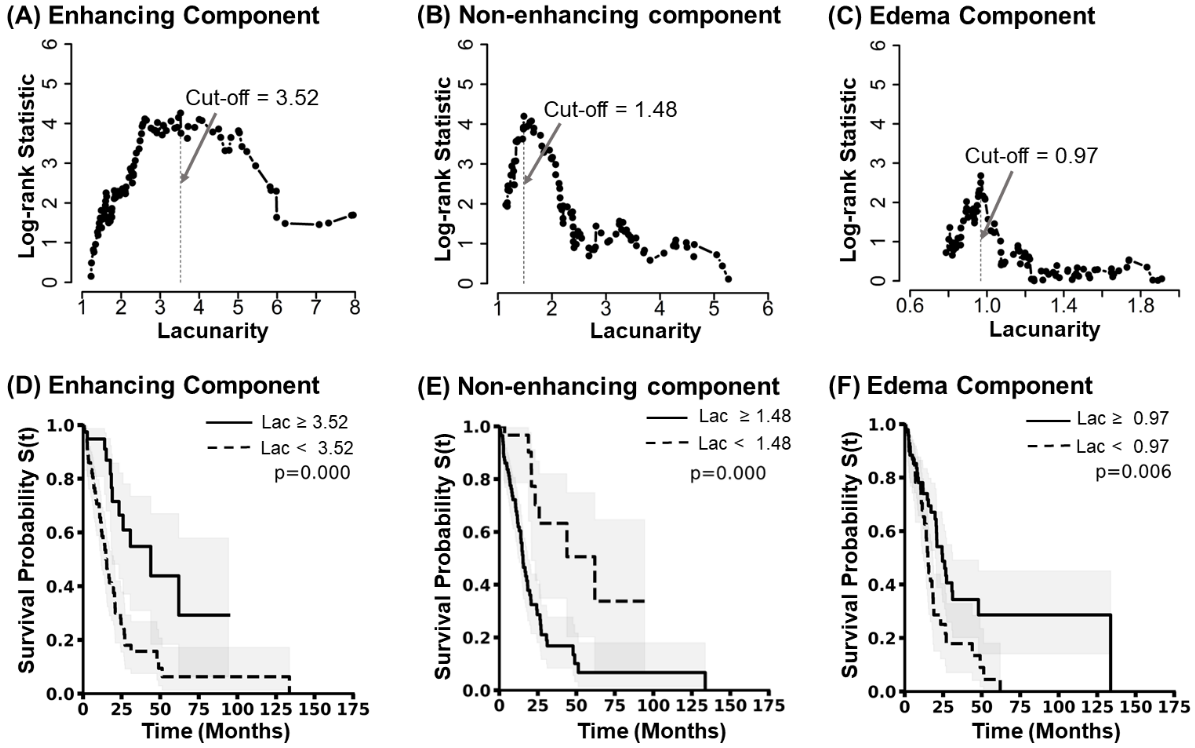


**Supplementary Figure S7: Lacunarity based determination of overall patient Survival**. Optimal cutoff values for the lacunarity of (A) enhancing, (B) nonenhancing, and (C) edema subcomponents were determined using MaxStat and log-rank tests. Kaplan-Meier survival curves between the two groups formed using the optimal cutoff values of The Fractal Dimension of (D) enhancing and (E) nonenhancing components and (F) edema tumor subcomponents. One group has an FD cutoff greater than and equal to the cutoff value (solid line), and the other has an FD less than the cutoff value (dashed line).
